# Oxytocin and vasotocin receptor variation sheds light into the evolution of human prosociality

**DOI:** 10.1101/460584

**Authors:** Constantina Theofanopoulou, Alejandro Andirkó, Cedric Boeckx, Erich D. Jarvis

**Affiliations:** Laboratory of Neurogenetics of Language, Rockefeller University, New York, NY, USA; Section of General Linguistics, Universitat de Barcelona; Universitat de Barcelona Institute for Complex Systems; ICREA; Howard Hughes Medical Institute, Chevy Chase, MD, USA

## Abstract

Modern human lifestyle strongly depends on complex social traits like empathy, tolerance and cooperation. These diverse facets of social cognition have been associated with variation in the oxytocin receptor (*OTR*) and its sister genes, the vasotocin/vasopressin receptors (*VTR1A*/*AVPR1A* and *AVPR1B/VTR1B*). Here, we compared the full genomic sequences of these receptors between modern humans, archaic humans, and 12 non-human primate species, and identified sites that show heterozygous variation in modern humans and archaic humans distinct from variation in other primates, and that have associated literature. We performed variant clustering, pathogenicity prediction, regulation, linkage disequilibrium frequency and selection analyses on data in different modern-human populations. We found five sites with modern human specific variation, where the modern human allele is the major allele in the global population (*OTR*: rs1042778, rs237885, rs6770632; *VTR1A*: rs10877969; *VTR1B*: rs33985287). Among them, the *OTR*-rs6770632 was predicted to be the most functional. We found two sites where alleles (*OTR:* rs59190448 and rs237888)^1^ present only in modern humans and archaic humans are under positive selection in modern humans, with rs237888 predicted to be a highly functional site. We identified three sites of convergent evolution between modern humans and bonobos (*OTR*: rs2228485 and rs237897; *VTR1A*: rs1042615), with *OTR*-rs2228485 ranking very highly in terms of functionality and being under balancing selection in modern humans. Our findings shed light on evolutionary questions of modern human and hominid prosociality, as well as on similarities in the social behavior between modern humans and bonobos.

## Introduction

Modern humans are characterized by prosociality, a broad term that encompasses intraspecies empathy, social tolerance, cooperation and altruism. While our closest living relatives, the chimpanzees (*Pan Troglodytes*) and the bonobos (*Pan Paniscus*), live in highly organized social groups as well, present-day humans’ social networks are larger and denser, powered by a complex social cognitive machinery^2^. Modern humans are also characterized by great intrasocial compassion and display a clear tendency to act in concert, to the extent that our species has been labeled as ‘ultra-social’^3^.

Less is known about the sociality of our extinct human relatives, the Neanderthals and the Denisovans. Paleogenomic studies show that both Neanderthal and Denisovan populations endured a population size bottleneck^4^. In addition to this, high inbreeding rates evident in both the Altai Neanderthal^5^ and the Denisovan^6^ genomes, as well as lower genetic diversity in Neanderthals compared to modern humans^7^ suggest that their social groups were most likely much smaller than ours. According to Rogers and colleagues^8^, Neanderthals were deeply subdivided into small isolated population groups with scarce contact between them, which may be associated with a social profile distinct from *Homo sapiens.*

A wide range of studies have highlighted a role of oxytocin and vasopressin/vasotocin receptor genes in social behaviors^9^ and associated disorders, such as in Autism Spectrum Disorders (ASD)^10,11^. These include *OTR (*a.k.a. *OXTR)*, *VTR1A (*a.k.a. *AVPR1A)* and *VTR1B (*a.k.a. *AVPR1B*), following the nomenclature proposed in^12^. These findings have led some researchers, most prominently Hare^13^, to ascribe to these genes a key role in the emergence of human ultra-social behavior.

In this study, we searched for sequence variation in the *OTR-VTR1* family of genes (specifically, *OTR, VTR1A* and *VTR1B)* that could shed light on the evolution of prosociality in modern humans. Since fixed or nearly fixed changes have not been found so far in these genes in comparisons between MH and archaic humans (AH) or non-human primates (NHP) (chimpanzees and bonobos in particular)^5,14–16^, we sought to identify potential polymorphic heterozygous sites in MH not found in AH or/and NHP, and we report those sites that have been associated with social features/deficits in the literature. We identified 5 sites with modern human specific variation, where the modern human alleles are the major alleles in the global population. Among them, sites in the *OTR* were in regulatory regions of the gene, active in the brain, and associated with prosocial behavior in modern humans. We also identified 3 convergent sites between modern humans and bonobos, a primate species that shows convergence of prosocial behaviors with humans, separate from chimpanzees. Some of the sites we report have been shown to be under balancing or positive selection in MH. Our findings shed light on the evolutionary questions on modern human and hominid prosociality, as well as on observations on similarities in the social behavior of MH and bonobos.

## Results

### Variation pattern analyses and associations with sociality

We performed alignments of the *OTR*, *VTR1A* and *VTR1B* genomic sequences (exon, introns, and surrounding regulatory regions 600 bp upstream and downstream) between seven high-coverage present-day human genomes (i.e. Modern Humans; MH), 14 Neanderthal genomes^5,17,18,19,20,21,22^ and a Denisovan^6^ genome (i.e. archaic humans; AH), and multiple genomes of 3 non-human primate (NHP) species (chimpanzee, bonobo, macaque; Supplementary Table 1). We also included several *VTR1A*-microsatellites that have been associated with social-related phenotypes^24,25^: RS3-(CT)_4_TT(CT)_8_(GT)_24_; RS1-(GATA)_14_; GT_25_; and AVR-(GT)_14_(GA)_13_(A)_8_. For the non-human primates, we additionally used available Single Nucleotide Variant (SNV)-data from bonobo, chimpanzee, and macaque populations (~30 to ~1000 individuals/species; **Supplementary Table 2**; Methods) to account for variation in non-human primates. We inferred the ancestral state of the sites we identified using Ortheus multialignments of 12 NHP species (Supplementary Note 1).

From the total number of Single Nucleotide Polymorphisms (SNPs) found on the *OTR* (5409 SNPs), *VTR1A* (2340 SNPs) and *VTR1B* (3031 SNPs) in the NCBI Variation Viewer (National Center for Biotechnology Information) in MH (GRCh38.p12 reference genome), only 79 had associated literature findings: *OTR* (55), *VTR1A* (7) and *VTR1B* (17). We excluded from our analysis those studies that review evidence from original studies, and we ended up with 61, consisting of 42 SNPs (285 studies) for *OTR*, 8 SNPs (24 studies) for *VTR1A*, and 11 SNPs (14 studies) for *VTR1B* (**Supplementary Tables 3, 4** and **5**). We classified these associations based on whether they were strongly associated to sociality (e.g. ‘prosociality’, ‘social cognition’), possibly related to sociality (e.g. ‘depression’, ‘Attention Deficit Hyperactivity Disorder’), or not related to sociality (e.g. ‘diabetes’). We then calculated the percentage of the associations for each site, and then averaged all the percentages for each category relative to the total number of the SNPs.

MH variant sites in *OTR* had the highest association with sociality (**Supplementary Table 3**): 72% of the findings were sociality-related (e.g. ‘empathy’, ‘social temperament’, ‘face recognition’); 14% possibly related to sociality (e.g. ‘obsessive compulsion disorder’, depression); while the remaining 14% was not related to sociality (e.g. ‘bulimia’, ‘overeating’, ‘diabetes’). These differences were significant (Chi-squared test; P < 0.0001). In contrast, variants in *VTR1A* had a weaker association, with 27% being related to sociality (e.g. ‘moral judgement’, ‘aggression’, ‘autism’), 6% possibly related to sociality (e.g. ‘susceptibility for panic disorder’), and 68% to other phenotypes (e.g. ‘nicotine dependence’, ‘heroin addiction’, ‘hypertension’; **Supplementary Table 4**; Chi-squared test; P = 0.0049). Concerning *VTR1B*, 50% of the findings were related to sociality (e.g. ‘emotional empathy’, ‘aggression’), 32% possibly related to sociality (‘bipolar disorder’, ‘psychotic features’), and 17% were not (e.g. ‘elevated body mass index’; **Supplementary Table 5**). Unlike the other two genes, these values for *VTR1B* were not significantly different between groups (Chi-squared test; P = 0.3417). Lastly, the difference between the proportions of the studies related to sociality were significant for *OTR* versus *VTR1A* (Chi-squared test; P < 0.0001), but not significant for *OTR* versus *VTR1B* (Chi-squared test; P = 0.0771) and *VTR1A* versus *VTR1B* (Chi-squared test; P = 0.1836; **Supplementary Table 6**). We interpret these findings as evidence for a higher functional association to sociality for *OTR* over *VTR1A* and *VTR1B*, although we are aware it could be a result of publication bias favoring sociality studies with *OTR.*

In total, of the 61 sites analyzed, 29 had associations with sociality (**Supplementary Tables 7-9**), while simultaneously showed a variation pattern found only in MH, in AH and MH, or in MH and a NHP species: 19 *OTR*-SNPs, 7 *VTR1A*-SNPs and 3 *VTR1B*-SNPs. Based on their variation patterns in MH, AH and NHP, we classified the 29 sites into five categories (**Figure 1**): 1) Modern Human Unique (MHU) for sites where the heterozyogous allelic variation is unique to MH (e.g. T/A in MH and A/A in all other species); 2) Modern Human Specific (MHS) for sites where MH have a specific variation pattern (e.g. T/A) different from that in other species (e.g. C/A); 3) Homo Unique (HU) for sites that are variant in both MH and AH in the same fashion (e.g. T/A), while the rest of the NHP are invariant (e.g. A/A) or have a different variant (e.g. C/A); 4) Homo Specific (HS) for sites that are variant in MH (e.g. T/A), and invariant in AH (e.g. T/T), different from all NHP species (e.g. A/A); And 5) MH-NHP for sites with convergent variation between MH (e.g. T/A) and another NHP (e.g. T/A in bonobo), where other species/populations are invariant for the ancestral allele (e.g. A/A). Of these 29 sites associated with sociality, most (13) were MHU in *OTR*, while the 3 in *VTR1B* were MHS. The others were in a mixture of genes, with 4 HS sites in *OTR* and 1 *VTR1A*; 2 HU sites in *OTR* and 1 in *VTR1A*; and 3 convergent sites between MH and bonobos in *OTR* and 1 in *VTR1A* (**Figure 1, Table 1**).

**Figure 1:**
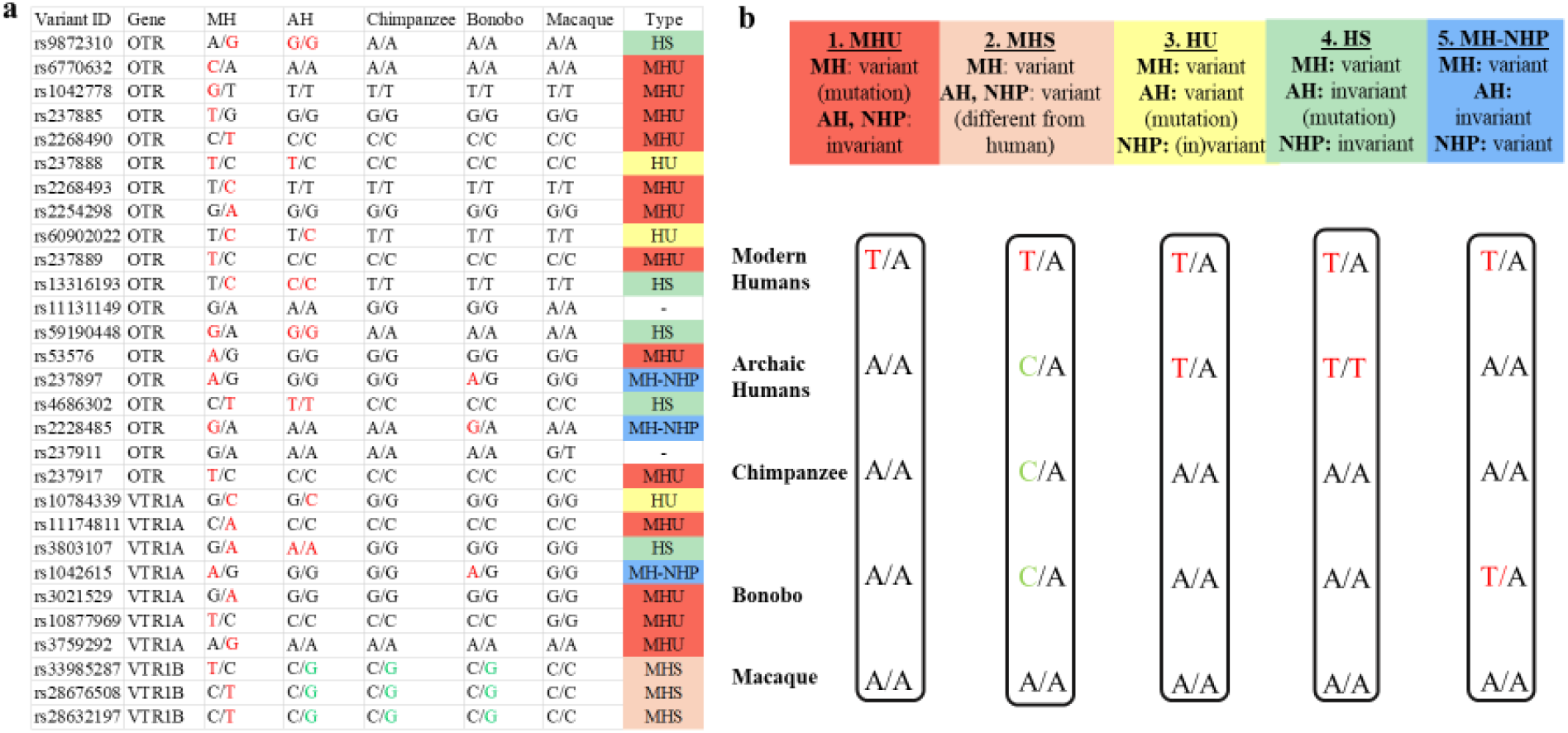
Variant clustering. **a**, For each variant site in *OTR, VTR1A* and *VTR1B*, we list the variation pattern found in MH (Modern Humans), AH (Archaic humans: Neanderthals and Denisovans), Chimpanzees, Bonobos and Macaques. In the column ‘Type’ we classify each allele on its assigned Type-category. Alleles that were key for each classification are in red font. **b**, A schematic representation of the four classification types. Shown in the columns are examples of each type. More details described for each type are in the main text results.

**Table 1:**
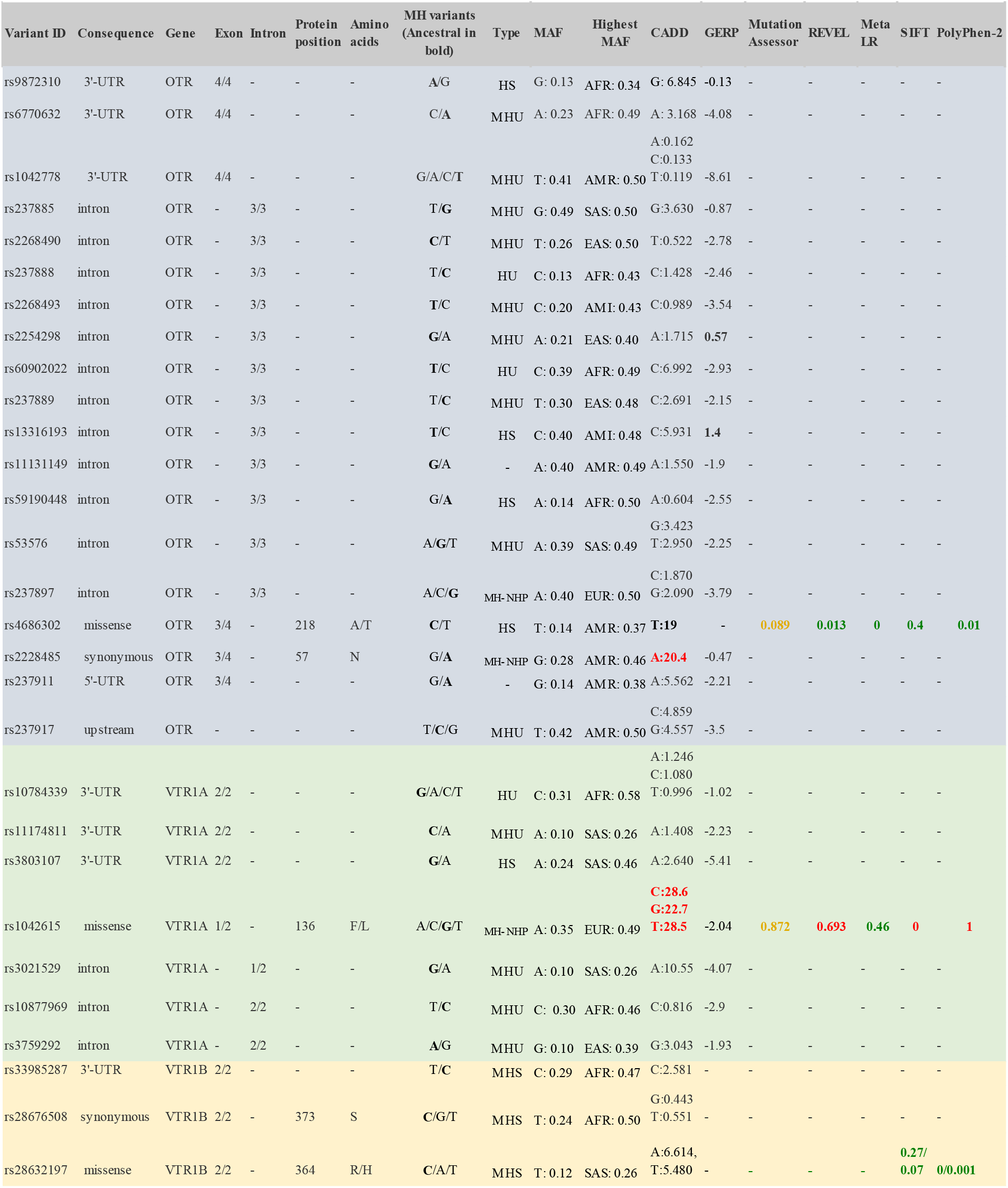
List of identified variants in Modern humans and pathogenicity prediction scores. For each variant, we have listed where it is found in the genetic sequence (intronic, 3′-UTR) and if in the exon, whether it gives rise to a missense or a synonymous mutation (‘Consequence’); on which specific ‘Exon’ or ‘Intron’ each site is located; on which ‘Protein position’ it is located, if exonic, and what ‘Amino acids’ it gives rise to; all MH (Modern human) alleles found on this site, with the ancestral being in bold; the ‘Type’ of each site based on our 5 categories (MHU: Modern Human Unique; MHS: Modern Human Specific; HU: Homo Unique; HS: Homo Specific; MH-NHP: Modern Human-Non-Human Primate); the Major Allele Frequency (MAF); the Highest Minor Allele Frequency (MAF) in a specific population from the 1000 Genomes or the gnomAD project (AFR: African, AMR: American, EAS: East Asian, SAS: South Asian, AMI: Amish); CADD (Combined Annotation Dependent Depletion) and GERP (Genomic Evolutionary Rate Profiling), Mutation Assessor, REVEL (Rare Exome Variant Ensemble Learner), MetaLR (Meta Logistic Regression), SIFT (Sorting Tolerant From Intolerant) and PolyPhen-2 scores. Scores in red: deleterious; scores in yellow: neutral; scores in green: benign.

Ancestral allele and functional association analyses revealed that in almost all cases, the ancestral and the MHU, MHS, HU or HS alleles showed mixed associations with sociality, i.e., the same allele was associated with both ‘prosocial’ and ‘antisocial’ features in the literature. In other words, we did not find a general trend where the ancestral allele would be associated with ‘antisocial’ features, and the new allele with ‘prosocial’ ones, or the other way around (**Supplementary Tables 7-9**). For example, the MHU *OTR*-rs1042778(G), which is highly present in MH populations, has been associated with ASD^26–28^, aggression in males, but also with prosocial fund allocations^29^ and high scores in altruistic and comforting behaviors^30^ (**Supplementary Table 7**). The ancestral T allele has been associated with diminished parental care^31^ and panic/aggressive behaviors^28^ on the one hand, but also more likely recovery from low maternal emotional warmth on the other^32^. A similar trend of mixed social associations for each variant, or absence of evidence for one of the allelic variants, was found for the rest of the sites (**Supplementary Table 7-9**).

### Pathogenicity prediction analyses

Most of these sites were located in regulatory regions of the genes, but some in exons giving rise to missense (*OTR*-rs4686302 [HS], *VTR1B*-rs28632197 [MHS] and *VTR1A*-rs1042615 [MH-NHP convergence]) or synonymous mutations (*VTR1A*-rs2228485 [MH-NHP convergence] and *VTR1B*-rs28676508 [MHS]) (**Table 1**). We performed pathogenicity prediction analyses using genome wide scores of conservation that predict deleteriousness of mutations (CADD^33^ [Combined Annotation Dependent Depletion] and GERP^34^ [Genomic Evolutionary Rate Profiling]), and algorithms that evaluate the impact of missense variants (Mutation Assessor^35^, REVEL^36^ [Rare Exome Variant Ensemble Learner], MetaLR^37^ [Meta Logistic Regression], SIFT^38^ [Sorting Tolerant From Intolerant] and PolyPhen-2^39^) (**Table 1**). CADD scores that predict variant deleteriousness showed that MH-NHP *OTR*-rs2228485(A) and *VTR1A*-rs1042615(G, C, T) alleles are likely deleterious (i.e. scores >20 predict deleteriousness), while HS *OTR*-rs4686302(T) also scored very close to this value (19; **Table 1**). Scores for most other sites ranged from ~0 to 6. GERP tests showed positive evolutionary rate scores (as defined in Methods) only for MHU *OTR*-rs2254298 and HS *OTR*-rs13316193 (**Table 1**), which represent highly conserved sites, that are likely to be functional. Combined REVEL, MetaLR, SIFT and PolyPhen-2 scores classified HS *OTR*-rs4686302 as more likely benign, with only the Mutation Assessor classifying it as likely deleterious (**Table 1**). MH-NHP *VTR1A*-rs1042615 was ranked by most of the aforementioned analyses as deleterious or likely deleterious, with only MetaLR ranking it as benign. SIFT and PolyPhen-2 tests categorized MHS *VTR1B*-rs28632197 as benign. These findings suggest that despite their deleterious potential, these sites have been maintained in the modern human population but not in other primates, perhaps due to selection differences in social behavior.

### Regulation analyses

We performed regulation analyses using RegulomeDB^40^ and the Ensembl regulation data resources^41^. RegulomeDB make use of large datasets, including ChIP-seq, FAIRE-seq and DNase I hypersensitive information for a variety of important regulatory factors across a diverse set of cell types; this includes chromatin state information for over 100 cell types, transcription factor binding sites (including Position-Weight Matrix and DNase Footprinting), and expression quantitative trait loci (eQTL) information allowing the association of distal sites with gene promoters. The Ensembl resources provide a functional annotation of the regulatory elements in the human genome, including data on epigenetic marks, transcription factor binding and DNA methylation, as well as microarray probe mappings.

Our RegulomeDB analyses ranked highly the MHU *OTR*-rs237889 variant site as lying in a functional genomic location (rank 1f; **Table 2**). Consistent with these findings, this same site is found in a genomic location with peak Chip-seq and DNase-seq signal for open chromatin in the brain and other cell types (**Figure 2**). This site is also located in an eQTL affecting expression of *CAMK1* and maps to the binding sites of the *EGR2* and *ZFHX2* transcription factors (**Table 2**). The second most highly ranked site was the MH-NHP *OTR*-rs2228485 allele (rank 2b). ChIP-seq, DNase-seq and FAIRE-seq data indicate that this site is in a genomic region active in the brain, and in particular the dopaminergic substantia nigra neurons and one of its projection targets, the caudate nucleus of the striatum, as well as the hippocampus, other parts of the temporal lobe, the cingulate and angular gyri, and the middle frontal area 46 (**Table 2**). This site is implicated in binding with *CTCF*, *EZH2*, *POLR2A*, *RAD21* and *IKZF1* transcription factors, but with the SNP potentially altering the binding for *PAX6* and *ZSCAN4C* (**Table 2**).

**Table 2:**
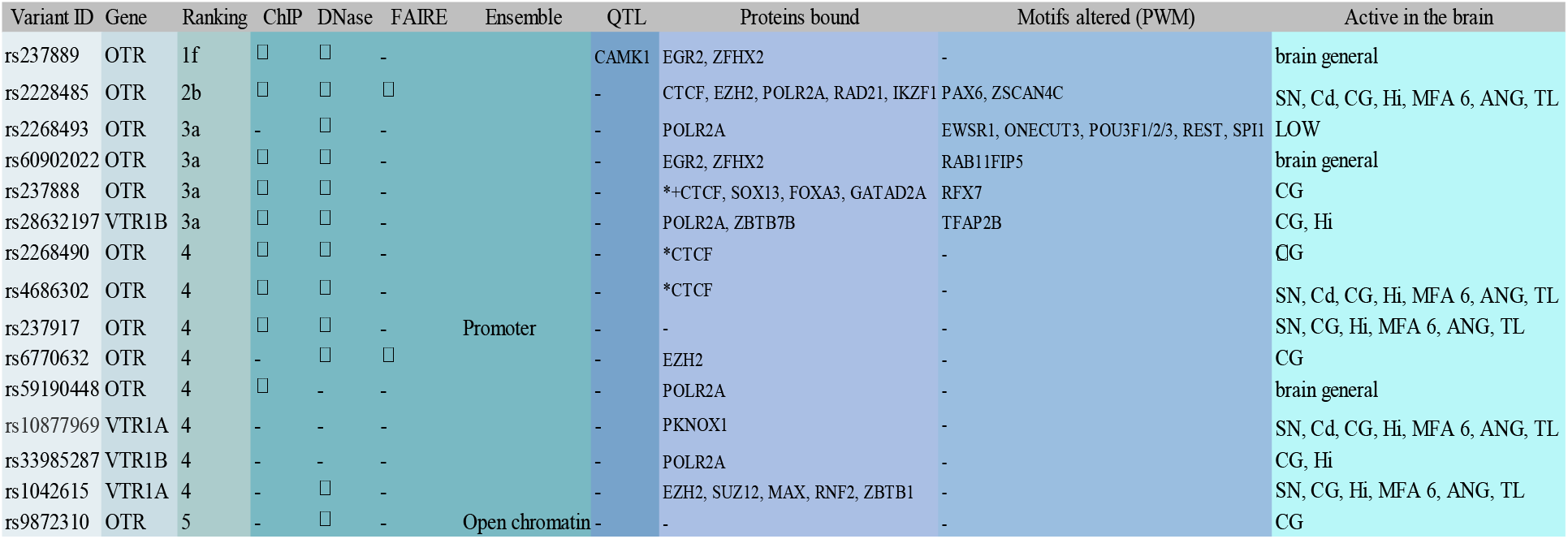
RegulomeDB high-ranked results and Ensembl regulation results. For each site, we list the gene they are found in, ranking scores from RegulomeDB, if they are found at open chromatin regions based on Chip-seq, DNase-seq or FAIRE-seq (IZ) data from Ensembl regulation database. This includes sites found on promoter or open chromatin regions, expression quantitative trait loci (eQTL) regions, proteins bound (* for those mentioned only in Ensembl, *+ for those supported from both RegulomeDB and Ensembl data. All the rest are come from RegulomeDB), PWM (positional weight matrix), whether they are active in the brain (e.g. in chromatin with evidence for strong transcription in the brain) and in which brain regions (SN: Substantia nigra; Cd: Caudate nucleus; CG: Cingular gyrus; Hi: Hippocampus; MFA 46: Middle Frontal Area 46; ANG: Angular Gyrus; TL: temporal lobe).

**Figure 2:**
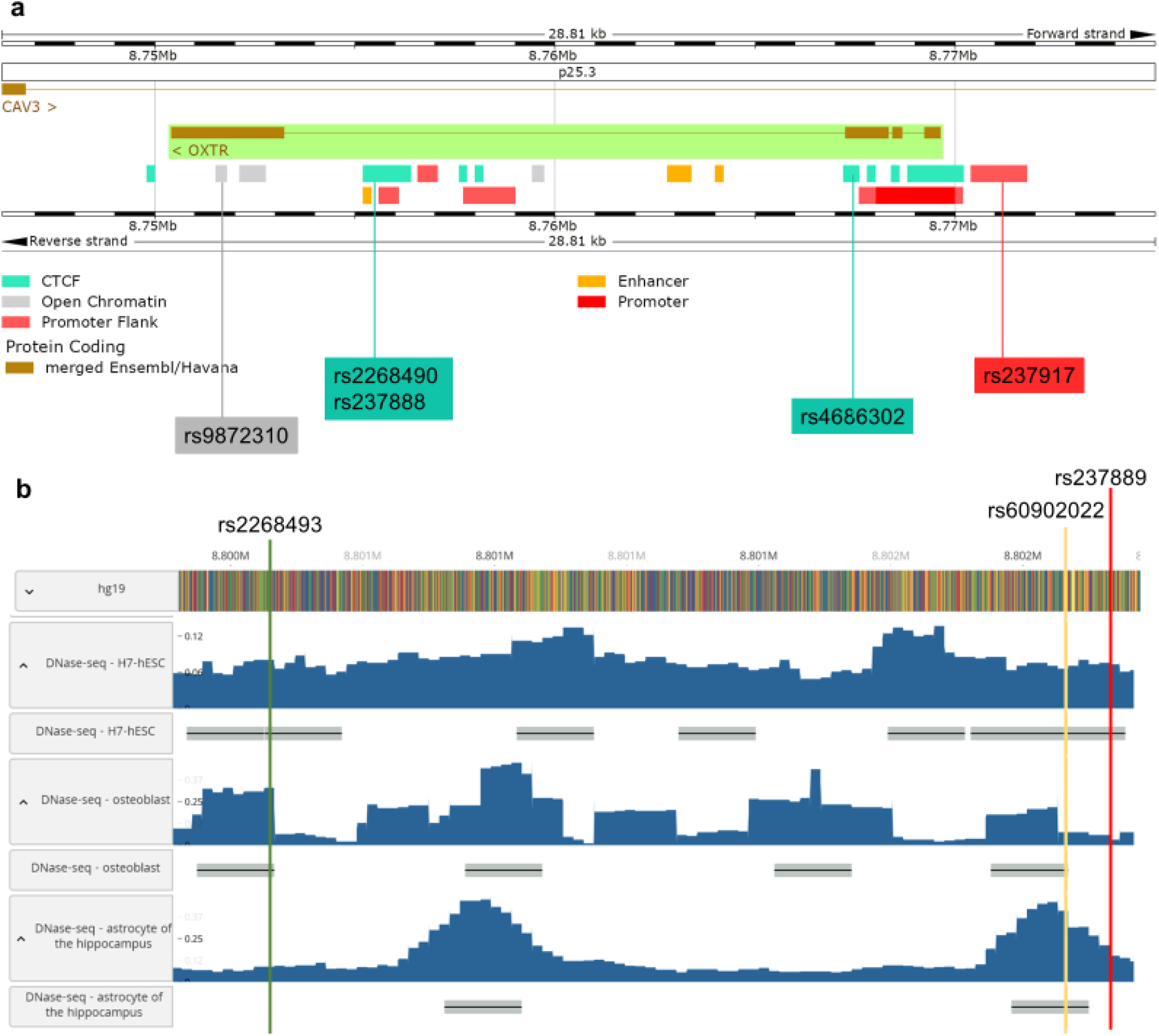
Ensembl regulation and RegulomeDB results. **a**, Ensembl regulation data resources for *OTR* (chromosome 3), showing the *CTCF* (transcription factor) binding domain (green), the open chromatin regions (grey), the promoter regions (red) and the enhancer regions (yellow). We highlighted SNP sites analyzed in this study. **b**, RegulomeDB browser (chromosome 3: 8,800-8,802 Mb, *OTR* intron) showing open chromatin regions (blue peaks) for 3 sites identified in this study. Shown are DNase sequencing data from undifferentiated embryonic stem cells (DNase-seq-H7-hESC), from osteoblasts, and from astrocytes from the hippocampus.

MH-NHP *OTR*-rs2228485, MHU *OTR*-rs2268493, HU *OTR*-rs60902022, HU *OTR*-rs237888 and MHS *VTR1B*-rs28632197 were all were less likely to affect regulation (rank 3a), but still had open chromatin regions that either fall within the binding site of different proteins, or alter gene motifs, or both, and showed activity in brain regions like the cingulate gyrus and the hippocampus (**Table 2; Figures 2a, 2b**). The remaining sites (*OTR*: rs2268490, rs4686302, rs4686302, rs237917, rs6770632, rs59190448 and rs9872310; *VTR1A*: rs10877969 and rs1042615; *VTR1B*: rs33985287) did not rank highly in changing regulation according to the RegulomeDB scores (ranks 4-6), but were also found in open chromatin or promoter regions (according to Ensembl or RegulomeDBdata), or in a transcription factor binding domain (**Figure 2a**), and all showed activity in the brain. A total of 8 of these 15 sites are located within the *CTCF* and the *POLR2A* transcription factor binding motifs (**Table 2; Supplementary Table 10**). These findings indicate that variant sites associated with sociality would cause variant brain gene expression among the MH population.

### Linkage disequilibrium analysis

We ran Linkage Disequilibrium analysis (using LDmatrix^42^) to test if any of the sites we identified are co-inherited, using data from all populations included in the 1000 Genomes Project and a cut off of R²> 0.8. This analysis revealed four haplotype blocks: one containing co-inheritances among rs60902022, rs13316193 and rs11131149 in *OTR*; a second between rs10784339 and rs10877969 in *VTR1A*; a third between rs11174811 and rs3021529 also in *VTR1A*; and a fourth between rs33985287 and rs28676508 in *VTR1B* (**Figure 3, Supplementary Table 11**). This suggests that the combinations of these variant alleles might act in concert to shape functional differences that end up affecting phenotypes controlled by these genes.

**Figure 3:**
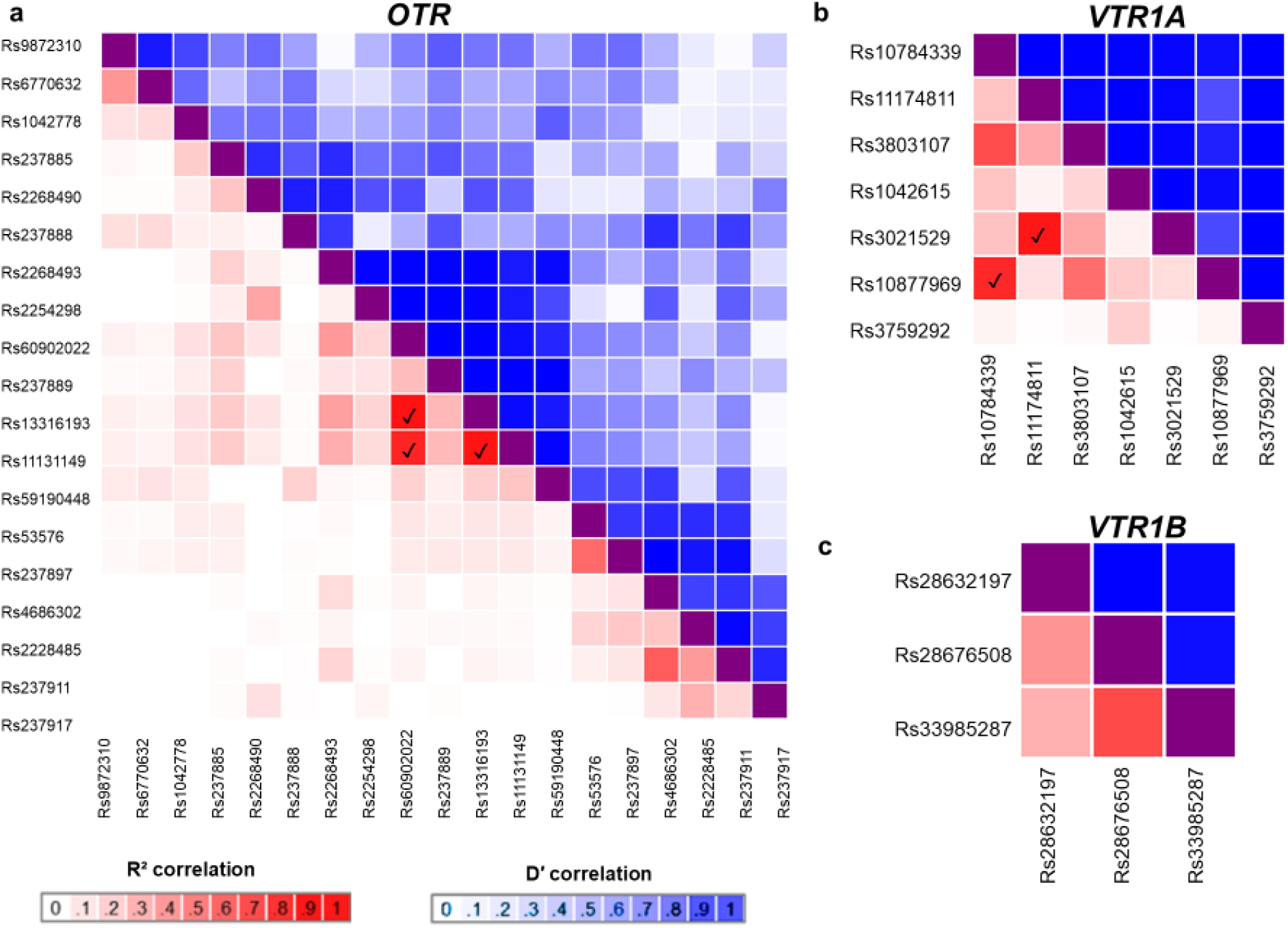
Linkage Disequilibrium analysis. **a**, *OTR* heatmap matrix of pairwise linkage disequilibrium statistics. Pairwise statistics are colored based on their R² and D’ significance scores, with the darkest shades of red being the most significant R² score, and white the least, and the darkest shades of blue being the highest D’ scores, and white the least. IZ, pairwise R²> 0.8, our threshold for a significant co-inheritance of the variant alleles. The same coloring and symbols were used in the rest of the heatmaps. **b**, *VTR1A* heatmap matrix of pairwise linkage disequilibrium statistics. **c**, *VTR1B* heatmap matrix of pairwise linkage disequilibrium statistics.

### Population data and selection analysis

We further analyzed the human allele frequency data for these sites (1000 Genomes Project^43^ Phase 3 and gnomAD^44^ v2.1.1 and v3) and the literature for selection on these sites (**Table 1, Supplementary Table 12, Extended Data Table 1**). We used the supplementary information in Schaschl et al. 2015^1^, and filtered the results of their FDIST, Bayescan and extended Lewontin and Krakauer (FLK) tests, reporting here only those on our identified SNPs with significant values (p or q values◻<◻0.05) (**Extended Data Table 1**). For bonobos, we also analyzed the data from 13 bonobos by Kovalaskas and colleagues^45^, where strong positive selection signals on *OTR* and *VTR1A* were identified.

We found four MHU and one MHS case where the new allele is currently the major allele in MH populations: (MHU: *OTR*-rs6770632(C), rs1042778(G), rs237885(T); MHS: *VTR1A*-rs10877969(T); *VTR1B*-rs33985287(T); **Table 1**). Several of these sites have a high derived allele frequency of ≥70% globally (*OTR*-rs6770632(C), *VTR1A*-rs10877969(T), *VTR1B*-rs33985287(T)), while *OTR*-rs1042778(G) and *OTR*-rs237885(T) showed lower frequencies (59% and 51%, respectively). Similarly, we found one HU and one HS case, where the new hominin (AH or MH) allele is the major allele in the global MH population (HU: *OTR*-rs59190448(G); HS: *OTR-*rs237888(T); **Table 1**). Both sites show derived allele frequencies of >85%, and they show signs of positive selection. Specifically, based on our filtering of the results in^1^, all three approaches (FDIST, BayeScan and FLK) identified the *OTR*-rs59190448(G) to be strongly under positive directional selection (FDIST p=0.0004, BayeScan q=0.0004, F.LK p=0.001), while only the FLK test (p=0.011) detected the *OTR-*rs237888 to be under positive selection (**Extended Data Table 1**). Interestingly, both the *OTR*-rs237888(T) and the *OTR*-rs59190448(G) are present in the majority of the AH genomes (6/8 for *OTR*-rs237888(T) and 5/7 for *OTR*-rs59190448(G); **Supplementary Table 1**); although, we acknowledge that the genomes of a small number of AH individuals currently available might not be representative of the general archaic population. These results, though tentative, show a similar frequency trend in AH and MH.

Several sites where we encountered a new allele either in AH or in MH are currently the minor allele in MH populations, with some of them being present at low percentages in the global population (<25%) (*OTR*-rs9872310, rs2268493, rs2254298, rs4686302, rs237911; *VTR1A*-rs11174811, rs3803107, rs3021529, rs3759292; and *VTR1B*-rs28676508, rs28632197). Our filtering of the results in^1^ showed that some of them are under balancing selection: *OTR*-rs237911 (FDIST, p= 0.0485), *VTR1A*-rs11174811 (FDIST, p=0; BayeScan, q=0.0336) and *VTR1A*-rs3021529 (FDIST, p= 0.0001; BayeScan q= 0.0266); only the *VTR1A*-rs3759292 was found to be under positive selection (F.LK, p=0.001; **Extended Data Table 1**). Interestingly, *VTR1A*-rs3759292 is also in a region that is found deleted in 0.3% of the macaques sampled on that site (739; **Supplementary Table 2**), and according to Donaldson and colleagues^25^, who found sites with both variation and deletion events in the MH and NHP *VTR1A* microsatellite, such regions might have influenced sociobehavioral traits during primate evolution. MH and bonobos showed convergent variation on three other sites (*OTR*-rs2228485 and rs237897; *VTR1A*-rs1042615). The *OTR*-rs2228485 allele was present in all the 13 bonobos of one population^46^ (87% of our total sample of 15 bonobos), but the alternate allele was present in the two reference genomes (13%). Although in all these sites, the convergent allele was not the major allele in the global MH population, in some MH populations it is present at ~50% (**Table 1, Supplementary Table 12**). In particular, the convergent *OTR*-rs2228485 G-allele that was found at 87% among the bonobo population, at 28% in the global MH population, while at 46% in the American population. *OTR*-rs237897(A) was found in 40% of the MH global population (50% in Europeans) and 13% in bonobos. *VTR1A*-rs1042615(A) reached 35% in the global population (49% in Europeans), and 87% in bonobos. Of these sites, only the *OTR*-rs2228485 was found to show signs of balancing selection (FDIST, p=0.0149) in MH. None of these convergent sites were among the specific sites of strong positive selection identified in bonobos^45^, but several of these sites were located in close vicinity to our identified sites in the *OTR* and *VTR1A* (e.g. one site was found at a ~14kb distance from *VTR1A*-rs1042615). These findings suggest a correlation of several sites associated with sociality also showing convergent changes in modern humans and bonobos, not found in chimpanzees and other non-human primates examined.

## Discussion

In this study, we find multiple cases of allelic heterozygosity variation in the *OTR*, *VTR1A* and *VTR1B* genes that is specific to MH or AH and MH, where one allele is unique to them among primates and the other shared with other primates. Variants in all three genes have been associated with sociality, although more so for *OTR* than *VTR1A* and *VTR1B*. As it is known that heterozygous variants can have different impacts on genes and traits than their homozygous counterparts, such as the protective affect for sickle cell anemia^47^, we suggest that these variants could have a synergistic heterozygous impact on the evolution of social behavior in hominids. However, because of the population variation, we believe that it may not be a specific allele that is important for increased sociality, but the combination of heterozygous alleles among the population that may influence the interactions differently than a population that were purely homozygous. SNPs can act as general risk factors, and not as specific modifier alleles in certain behavioral or biological contexts (i.e. disorder specific effects)^48^. Certain alleles associated with more social deficits in one disorder (e.g. ASD) had either no effect, or conferred a benefit with respect to social abilities in another disorder (e.g. ADHD). Additionally, the impact of the alleles of a certain variant site is different depending on environmental factors, because environmental processes may be differentially accentuated through hormone signaling. We thus focus our interpretations on the functional and frequency analyses.

We consider that of the MHU and MHS SNPs, only those five where the new allele is the major allele in MH populations could be responsible for beneficial changes in MH relevant to sociality (*OTR*-rs6770632, rs1042778, rs237885; *VTR1A*-rs10877969; *VTR1B*-rs33985287), especially those found at ≥70% (*OTR*-rs6770632, *VTR1A*-rs10877969, *VTR1B*-rs33985287). *OTR*-rs6770632 is the only site among these three that falls in open chromatin regions active in the brain, and particularly in the cingulate gyrus, a brain region involved in social cognition^49^. Moreover, this site is in a genomic region containing the binding domain of *EZH2*, a gene that is involved in the etiology of memory impairments^50^, autism^51^ and Weaver syndrome, an overgrowth/intellectual disability syndrome^52^.

These MHS sites can shed light on several working hypotheses that account for MH prosociality. Among them, the ‘self-domestication hypothesis’^13,53–56^ posits that certain physiological and behavioral traits that MH share with domesticated animals support a significant turning point exclusive of MH on the prosocial continuum. Interestingly, Hare^13^ and Theofanopoulou^54^ have hypothesized that human self-domestication was likely facilitated by changes in the oxytocin/vasotocin system, based in part on studies showing that oxytocin and vasotocin receptors are under relaxed selective constraint in domesticated species^57^, and their gene expression and methylation patterns differ in wild versus domesticated species^58,59^. We also found that some of the variants found only in MH among primates (*VTR1A*-rs11174811 and *VTR1A*-rs3021529) show signs of relaxed selection.

Linkage disequilibrium analyses reveal more intricate relationships between alleles that might act in unison, likely conferring differential effects in specific MH populations. For example, the MHU *VTR1A*-rs10877969 variant co-inherited with the HS *VTR1A*-rs10784339, although not present in high percentages globally (31%), could be responsible for differences in the African population (58% in the African population). Similarly, other alleles that are found in low percentages in MH but showed highly likely functional results, like the *OTR*-rs237889 (30% in the global population; highest score (1f) in RegulomeDB analysis), might be responsible for characteristics of the Japanese population, where the MHS allele is more frequent (48%). According to Schaschl and colleagues^1^ who studied selection patterns of *OTR* and *VTR1A* SNPs in different human populations, the different population distribution of these alleles could influence the emergence of differences in socio-cultural practices on interpersonal proximity and touch, emotional suppression, social or emotional support, use of facial expression, inter-individual distance, and even susceptibility to ASD.

HU sites, where AH is homozygous with a new mutation not found in NHP, and MH are variant, can offer information on those SNPs that were likely present in AH and were selected for or purged out in the MH populations. *OTR*-rs59190448(G) and *OTR*-rs237888 are the only two sites where the HU allele ended up being positively selected in MH^1^, possibly resulting in beneficial phenotypes. With the open chromatin region for *OTR*-rs237888 also in the cingulate gyrus, we propose that this site along with the aforementioned *OTR-* rs59190448, could account for a more prosocial phenotype in the human lineage compared to our non-human primate ancestors at the time of the *Pan*-*Homo* lineage split^60,61^. Contrariwise, some of the other HU alleles that were found in open chromatin or conserved sites, have not been highly retained in the global MH population (*OTR*: rs9872310 [open chromatin], rs13316193 [conserved site], rs4686302 [open chromatin, mixed results on whether it is benign or deleterious]; *VTR1A:* rs3803107).

For the three sites of convergent evolution between MH and bonobos, *OTR*-rs2228485 was found to be under balancing selection in MH and is also one of the most highly ranked sites in our RegulomeDB analysis, pointing to putative high functionality. Functionality in this site with balancing selection might also contribute to phenotypic plasticity, as has been shown for other sites^62^. Only the *OTR*-rs2228485(G) and the *VTR1A*-rs1042615(A) were the major alleles in our bonobo sample, but in order to conclusively resolve the state of these sites in bonobos, larger population samples will be needed. Nonetheless, our findings on convergent changes for MH with bonobos could be insightful for understanding the posited similarities in prosociality between MH and bonobos, and the differences of both compared to chimpanzees^63,64^. Added to that, according to a recent study^45^, bonobos, since their split from chimpanzees, show strong positive selection signals on variants located in genes of the oxytocin and vasotocin pathway (including *OTR* and *VTR1A*), something they consider as evidence for reduced aggression and self-domestication in bonobos.

In conclusion, we found more evidence (5 sites in oxytocin and vasotocin receptors) pointing towards a prosocial shift in MH, and less (2 sites) supporting a similar shift in the hominin populations altogether. Although the fact that there are vastly more MH genomes currently available that might tip the balance towards MH specificity, the contrasting patterns obtained for oxytocin and vasotocin receptors suggest that our results cannot be fully reduced to the number of genomes available. Other receptors and neural ligands besides *OT* and *VT* with early changes in hominids, such as dopamine^61^ and β-endorphines^60^, are possible. When considering these hypotheses, one will need to consider that all of these ligands are known to interact^65^, so it could be that the early changes in *OTR* identified here formed part of a broader set of changes that set the stage for our prosocial profile. Our results and interpretations are also broadly compatible with information based on the fossil record, paleogenomic evidence^5–8^, and with behavioral differences between chimpanzees and bonobos^63,64^.

## Methods

### Variation pattern analyses

We retrieved the *OTR*, *VTR1A* and *VTR1B* DNA sequences from the following sources: the publicly available genomes of 14 Neanderthal individuals^5,17,18,19,20,21,22^ and a Denisovan^6^; seven high-coverage present-day human genomes (San(HGDP01036), Mbuti(HGDP00982), Karitiana(HGDP01015), Yoruba(HGDP00936), Dinka(DNK07), French(HGDP00533) and Han(HGDP00775), originally sequenced for^5^; 1000 Genomes project data Phase 3^43^ (as shown in Ensembl^66^); a chimpanzee (*Pan Troglodytes*) (CHIMP2.1.4/panTro4), bonobo (*Pan Paniscus*) (panpan1.1/panPan2) and rhesus macaque (*Macaca Mulatta*) (BMC Mmul_8.0.1/rheMac8).

We ran a multi-alignment between the modern human, archaic human, chimpanzee, bonobo and macaque gene sequences of *OTR*, *VTR1A* and *VTR1B*. Of the differences we found, we focused on those which are polymorphic in MH with literature associated with sociality or social deficits. We performed the alignments using the ‘Phylogenetic Context’ tool built in Ensembl^66^, tools in the Max Planck for Evolutionary Anthropology Ancient Genome Browser (https://bioinf.eva.mpg.de/jbrowse/), including MUSCLE^67^ and MView^68^ to generate the alignments, and Aliview^69^, Decipher for R^70^, Bedtools to edit and analyze them. We used all the genomic sequence of the genes we aligned, as provided in the standard layout of the files of the genomic sequences in the Ensembl database, namely with 600 bp upstream and downstream. We defined the genomic sequences in the same way when we extracted the gene sequences from the archaic genomes. The only additional regions we included ad hoc in our analysis that fall further out of 600bp window were those of several *VTR1A*-microsatellites that have been associated with social-related phenotypes^24,25^ (RS3-(CT)_4_TT(CT)_8_(GT)_24_, RS1-(GATA)_14_, GT_25_, and AVR-(GT)_14_(GA)_13_(A)_8_).

We then aligned these sites with the rest of the available Neanderthal genomes (AH: Spy^18^ (2 individuals), Goyet^19^, Les Cottés^20^ (5 individuals), Hohlenstein-Stadel^21^, Scladina^21^, Chagyrskaya^23^) and additional NHP genomes, chimpanzee (*Pan Troglodytes*) (Pan_tro 3.0/ panTro5 and Clint_PTRv2/ panTro6), bonobo (*Pan Paniscus*) (Max-Planck Institute panpan1/panPan1) and Rhesus macaque (*Macaca Mulatta*) (Mmul_10/ rheMac10). Of the AH sites, we included only sequence reads that had a mapping quality of ≥25. Due to low-quality DNA for most of the ancient samples, and given the heterogeneity of procedures for data production in AH genome sequencing, we considered variant alleles only those present in at least two AH genomes, to avoid false positives due to sequencing errors. Following^71^, we filtered out the samples that strongly deviated from the remaining individuals in the dataset. This filtering of samples only affected two major alleles discussed in the study: the MH-bonobo convergent *VTR1A*-rs1042615(A), that is also found in the Denisovan sequence; and the MHS *OTR*-rs6770632(C) allele that is also found in the Altai Neanderthal sequence. All alleles we retrieved can be found in Supplementary Table 1.

To assess variation in NHP, we used Single Nucleotide Variant (SNV)-data from: bonobos (13 bonobos from^46^, 27 individuals with only *OTR* and *VTR1A* data from^72^, 113 individuals with only *VTR1A* data from^73^); chimpanzees (25 individuals from^46^, 35^72^ and 62^74^ individuals with only *VTR1A* and *OTR* data); and macaques (1,234 individuals from the mGAP database^75^ and dbSNP 150 in Ensembl^66^, where the mGAP data have been incorporated) (Supplementary Table 2). The SNV-data from bonobos and chimpanzees from^46^ were lifted over to the GRCh38/hg38 version of the human genome (*Homo sapiens*), while the macaque SNVs^75^ were lifted to the GRCh37/hg19 version of the human genome; they were lifted from different human references because these are where the different primate data sets were mapped to. We lastly used information from several other publicly available genomes of these species, since they were sequenced from different individuals (chimpanzee: Pan_tro 3.0/ panTro5 and Clint_PTRv2/ panTro6, bonobo: Max-Planck Institute panpan1/panPan1, and rhesus macaque: Mmul_10/ rheMac10).

### Ancestral allele inference

In order to infer the ancestral alleles of the identified sites, we aligned the MH sequences of *OTR*, *VTR1A* and *VTR1B* against 12 non-human primate species, and inferred the ancestral sequences through Ortheus^76^ using the ‘Phylogenetic Context’ tool built in Ensembl^66^. Ortheus is a probabilistic method that infers the ancestral allele, using a phylogenetic model, that incorporate gaps, to infer insertion and deletion events. Ancestral sequences are predicted for each node of the phylogenetic tree. Multiple alignments of all the sites we identified in 12 primate species, with the inferred ancestral sequences in each node, can be found in Supplementary Note 1. Although we found several other putative sites of convergent evolution between humans and other non-human primates (e.g. mouse lemur, olive baboon, gelada etc.; **Supplementary Note 1**), in absence of variation data in these species we can’t determine the reliability of this convergence as we have done with bonobo population data.

### Association studies analysis

We went through all the association studies identified in the NCBI Variation Viewer^77^ as of October 2020 (e.g. for *OTR*: https://www.ncbi.nlm.nih.gov/variation/view/?q=OXTR) for SNPs that have publications (Filters: ‘Variant type’ (: Single Nucleotide Polymorphism), and ‘Has publications’ (: Yes)), and report the association(s) identified for each SNP site (**Supplementary Tables 3, 4 and 5**). We then classified these associations on whether they were strongly related to sociality (e.g. ‘prosociality’, ‘social cognition’), ‘possibly related to sociality’ (e.g. ‘depression’, ‘Attention Deficit Hyperactivity Disorder’), or not (e.g. ‘diabetes’). We calculated the percentage of the associations for each site first, to then add all the percentages for each category together averaged out by the total number of the SNPs. Although there are more studies on the *OTR* SNPs in total, mostly due to replication bias, we normalized this factor, by calculating an association-percentage for each SNP first (whether it came from one or several studies), and then used the number of the SNPs to average out the final results (instead of the number of studies). We calculated p values between the percentages, using a Chi-squared test (n exact sample sizes, degrees of freedom and confidence intervals are noted in **Supplementary Tables 3, 4 and 5**). We lastly calculated the difference between the proportions of the studies related to sociality for each pair-wise comparison of the analyzed genes, using a Chi-squared test (n exact sample sizes, degrees of freedom and confidence intervals are noted in **Supplementary Table 6**). All the sites we report have studies that we classified as clearly related to sociality, with the exception of rs10784339 that has been associated with substance use disorders (**Supplementary Tables 7-9**). Although we classified those studies as not related to sociality, we included this site in the analysis, recognizing that there might be subjectivity in the classification of substance use disorders as being related to sociality or not.

For some sites where we identified variation patterns we wanted to study further in terms of disorders, we performed an additional exhaustive literature research in the National Center for Biotechnology Information^77^ (https://www.ncbi.nlm.nih.gov/pubmed/), SNPedia (http://snpedia.com) and Google Scholar (https://scholar.google.com/platforms) (as of October 2020) for the clinical significance of each one of the SNPs we identified in present-day human populations. On Supplementary Tables 10-12, we have included most of studies that report an association of these variants, a short description of their exact impact, as well as the trial sample of each study. We have not included review studies that recapitulate findings of original studies or studies showing negative results.

### Pathogenicity prediction analyses

We performed pathogenicity prediction analyses using the following seven tools:

1. CADD^33^ (Combined Annotation Dependent Depletion), which scores the predicted deleteriousness of SNVs and insertion/deletions variants in the human genome by integrating multiple annotations including conservation and functional information into one metric. In our analysis, we display scores above 20 as likely deleterious and scores below 20 as likely benign.
2. GERP^34^ (Genomic Evolutionary Rate Profiling), which identifies constrained loci in multiple sequence alignments by comparing the level of substitution observed to that expected if there was no functional constraint. Positive scores represent highly-conserved positions while negative scores represent highly-variable positions.
3. Mutation Assessor^35^, which predicts the functional impact of amino-acid substitutions in proteins using the evolutionary conservation of the affected amino acid in protein homologs. The rank score is between 0 and 1, with variants with higher scores being more likely to be deleterious.
4. REVEL^36^ (Rare Exome Variant Ensemble Learner), which is an ensemble method for predicting the pathogenicity of missense variants. It integrates scores from MutPred, FATHMM v2.3, VEST 3.0, PolyPhen-2, SIFT, PROVEAN, MutationAssessor, MutationTaster, LRT, GERP++, SiPhy, phyloP, and phastCons. Scores range from 0 to 1, and variants with higher scores are predicted to be more likely to be pathogenic. We considered scores above 0.5 as ‘likely disease causing’ and below 0.5 is ‘likely benign’.
5. MetaLR^37^ (Meta Logistic Regression), which uses logistic regression to integrate nine independent variant deleteriousness scores and allele frequency information to predict the deleteriousness of missense variants. Variants are classified as ‘tolerated’ or ‘damaging’ scores rank between 0 and 1, and variants with higher scores are more likely to be deleterious.
6. SIFT^38^ (Sorting Tolerant From Intolerant), which predicts whether an amino acid substitution is likely to affect protein function based on sequence homology and the physico-chemical similarity between the alternate amino acids. Scores < 0.05 are classified as ‘deleterious’ and all others as ‘tolerated’.
7. PolyPhen-2^39^, which predicts the effect of an amino acid substitution on the structure and function of a protein using sequence homology, Pfam annotations, 3D structures from PDB where available, and a number of other databases and tools (including DSSP, ncoils etc.). The PolyPhen-2 score represents the probability that a substitution is damaging, so values nearer 1 are more confidently predicted to be deleterious (note that this is opposite to SIFT).

### Regulation analyses

We performed regulation analyses using RegulomeDB^40^ and Ensembl regulation data resources^41^. RegulomeDB analysis make use of large datasets, including ChIP-seq, FAIRE-seq and DNase I hypersensitive information for a variety of important regulatory factors across a diverse set of cell types, chromatin state information across over 100 cell types, binding sites of transcription factors (including Position-Weight Matrix and DNase Footprinting), and expression quantitative trait loci (eQTL) information allowing the association of distal sites with gene promoters. The Ensembl resources provide a functional annotation of the regulatory elements in the human genome, including data on epigenetic marks, transcription factor binding and DNA methylation, as well as microarray probe mappings (**Table 2, Figure 2, Supplementary Table 10**).

RegulomeDB has developed a heuristic scoring system based on functional confidence of a variant. Category 1 variants are those that are known eQTLs for genes, and thus have been shown to be associated with expression. Within Category 1, subcategories indicate additional annotations from the most confident (1a, which has transcription factor (TF) binding, a motif for that TF, and a DNase footprint) to the least confident (1f, which has only TF binding or a DNase peak). According to Boyle and colleagues^40^ who developed the system: ‘*the additional categories represent analogous annotations to Category 1 but without eQTL data and, thus, no known direct effect on binding. Category 2(a–c) demonstrates direct evidence of binding through ChIP-seq and DNase with either a matched PWM (positional weight matrix) to the ChIP-seq factor or a DNase footprint. Category 3(a–b) is considered less confident in affecting binding due to a more incomplete set of evidence. These sites have ChIP-seq evidence and either a motif that matches the ChIP-seq data but no DNase evidence, or DNase evidence and any other motif. Finally, Categories 4–6 lack evidence of the variant actually disrupting the site of binding. These include DNase and ChIP-seq evidence (Category 4), DNase or ChIP-seq evidence (Category 5), or any single annotation not in the above categories (Category 6).*’ A list of the scores of all variants studied can be found in Supplementary Table 7.

### Linkage Disequilibrium analysis

We ran Linkage Disequilibrium analysis (using LDmatrix^42^) to test if any of the sites we identified get coinherited, using data from all population included in the 1000 Genomes Project and a cut off of R²= 0.8. LDMatrix (https://ldlink.nci.nih.gov/?tab=ldmatrix) creates an interactive heatmap matrix of pairwise linkage disequilibrium statistics. R² and D’ scores for all sites’ combinations can be found in Supplementary Table 8.

### Population data and selection analysis

We collected human allele frequency data for our identified sites using data from the 1000 Genomes Project^43^ (Phase 3) and gnomAD v2.1.1 and v3^44^. We listed the minor allele frequency (MAF) and the highest MAF in present day human populations in Table 1, and specific frequencies only from the 1000 Genomes Project in Supplementary Table 9. For bonobos, we also analyzed the data from 13 bonobos by Kovalaskas and colleagues^45^ (raw data provided by Sarah Kovalaskas), where strong positive selection signals on *OTR* and *VTR1A* were identified.

We also reviewed the literature for studies on selection signals of these sites in MH. We used the supplementary information found in Schaschl et al. 2015^1^, and filtered the results of their FDIST, Bayescan and extended Lewontin and Krakauer (FLK) tests, reporting only those on our identified SNPs with significant values (p or q values◻<◻0.05). In their study, Schaschl et al. 2015^1^ reported in the main text only those SNPs with p or q values◻<◻0.005. Schaschl et al. 2015^1^ analyzed 14 human populations (n◻=◻1092) from the 1000 Genomes database and aimed to identify SNPs under selection employing two tests that use Bayesian models to detect F_ST_ outliers (FDIST and Bayescan) and the FLK test that compares patterns of differences between allele frequencies in several populations relative to their expectation under neutral evolution (details for each analysis in the Methods of Schaschl et al. 2015^1^).

## Acknowledgements

We thank Thomas O’Rourke for comments on the manuscript. We also thank Evan E. Eichler for guidance on primate variation data.

## Author Contributions Statement

CTh conceptualized and designed the study, ran variation, pathogenicity prediction, regulation, human population and macaque data, and association analyses. AA ran variation, pathogenicity prediction, human, chimpanzee and bonobo variation analyses and reviewed the literature. EDJ and CB supervised the study.

## Data availability statement

All data generated or analyzed during this study are included in the Suppl. Info and Tables.

---

**Extended Table 1:** Selection analysis based on Schaschl et al. 2015^1^ results. Listed are the results from the FDIST, Bayescan and Lewontin and Krackauer extended tests of our identified SNPs found in the Schaschl et al. 2015^1^ Additional File. The results with significance based on our filtering (P or q value◻<◻0.05) are highlighted in green (: positive selection) or yellow (: balancing selection). In FDIST: *H_E_*, expected heterozygosity; *F_ST_*, locus-specific genetic divergence among human populations. In BayeScan (User Manual; http://cmpg.unibe.ch/software/BayeScan/files/BayeScan2.1_manual.pdf): log10(BF)◻= 1 indicate ‘strong’ evidence for selection and >1.5 indicate ‘very strong’ evidence for positive selection; q-val, the BayeScan analogue of a P value; αi < 0, balancing or purifying selection; αi = 0; diversifying selection (αi > 0); *F_ST_*, locus-specific genetic divergence among human populations. In FLK: Ht, estimated heterozygosity, Lewontion and Krackauer extended test statistics (F.LK) and P values for F.LK.*F_ST_* values differ due to different calculation algorithms.

| rsID | FDIST |  |  | BayeScan |  |  |  | FLK |  |  |
| --- | --- | --- | --- | --- | --- | --- | --- | --- | --- | --- |
| | $H_E$ | $F_{ST}$ | $P$ | log10(BF) | q-val | $\alpha_i$ | $F_{ST}$ | Ht | F.LK | $P$ |
| rs6770632 | 0.3567 | 0.1545 | 0.9551 | -0.3203 | 0.2942 | 0.2607 | 0.1131 | 0.423 | 3.988 | 0.136 |
| rs1042778 | 0.4776 | 0.1635 | 0.9851 | -1.2907 | 0.8133 | 0.0161 | 0.0891 | 0.467 | 0.142 | 0.932 |
| rs237885 | 0.5013 | 0.0555 | 0.2712 | 0.1385 | 0.1806 | -0.6004 | 0.0581 | 0.499 | 0 | 1 |
| rs237888 | 0.196 | 0.1925 | 0.798 | -0.6614 | 0.4348 | 0.1349 | 0.1009 | 0.267 | 8.993 | 0.011 |
| rs59190448 | 0.1825 | 0.3103 | 0.0004 | 3.3977 | 0.0004 | 1.447 | 0.2548 | 0.262 | 13.521 | 0.001 |
| rs10877969 | 0.366 | 0.1935 | 0.8974 | -1.3998 | 0.8156 | 0.0046 | 0.0738 | 0.377 | 2.584 | 0.275 |
| rs11174811 | 0.1467 | 0.0047 | 0 | 1.2742 | 0.0336 | -1.3077 | 0.0255 | 0.136 | 0.622 | 0.733 |
| rs3021529 | 0.1567 | 0.0057 | 0.0001 | 1.2035 | 0.0399 | -1.2604 | 0.0266 | 0.147 | 0.594 | 0.743 |
| rs3759292 | 0.1552 | 0.2003 | 0.1136 | -0.7408 | 0.3899 | 0.1121 | 0.0832 | 0.176 | 14.129 | 0.001 |
| rs237911 | 0.2718 | 0.0315 | 0.0485 | -1.1798 | 0.7611 | -0.0238 | 0.0864 | 0.266 | 2.701 | 0.259 |
| rs2228485 | 0.4217 | 0.0238 | 0.0149 | -1.1187 | 0.7285 | -0.0313 | 0.086 | 0.437 | 3.486 | 0.175 |

