## Supplementary note 1, Supp. tables 7-9 for "Oxytocin and vasotocin receptor variation sheds light into the evolution of human prosociality"

4 ICREA

5 Howard Hughes Medical Institute

^+^these authors contributed equally to this work

**Supplementary Information**

**Supplementary Note 1**

We list all the multialignments conducted on the OTR, VTR1A and VTR1B sites discussed in the study. Alignments were done using Ortheus^1^, as it is built in the Ensembl^2^ ‘Phylogenetic context’ tool, in order to infer the ancestral alleles of the identified sites. Ortheus is a probabilistic method for the inference of ancestor, whose main contribution is the use of a phylogenetic model incorporating gaps to infer insertion and deletion events. Ancestral sequences are predicted for each node of the phylogenetic tree that relates the sequences. Human (*Homo sapiens*; GRCh38.p13/hg38) sites were searched in Ensembl with their rsIDs, and alignments were made with human as reference. 12 non-human primate species were used to infer the ancestral states: Bonobo (*Pan paniscus*; panpan1.1), Chimpanzee (*Pan troglodytes*; Pan_tro_3.0), Gorilla (*Gorilla gorilla*; gorGor4), Orangutan (*Pongo abelii*; PPYG2), Gibbon (*Nomascus leucogenys*; Nleu_3.0), Vervet (*Chlorocebus sabaeus*; ChlSab1.1), Crab-eating macaque (*Macaca fascicularis*; Macaca_fascicularis_5.0), Macaque (*Macaca mulatta*; Mmul_10), Olive baboon (*Papio Anubis*; Panu_3.0), Gelada (*Theropithecus gelada*; Tgel_1.0), Marmoset (*Callithrix jacchus*; ASM275486v1) and Mouse Lemur (*Microcebus murinus*; Mmur_3.0).

```
OTR
rs2228485(G/A)

Human                  GCACACACGCGTTCCCGCTCA
Ancestral sequences 1  GCACACACGCATTCCCGCTCA
Bonobo                 NNNNNNNNNNNNNNNNNNNNN
Ancestral sequences 2  GCACACACGCATTCCCGCTCA
Chimpanzee             GCACACACGCATTCCCGCTCA
Ancestral sequences 3  GCACACACGCATTCCCGCTCA
Gorilla                GCACACACGCATTCCCGCTCA
Ancestral sequences 4  GCACACACGCATTCCCGCTCA
Orangutan              GCACACACGCATTCCCGCTCA
Ancestral sequences 5  GCACACACGCATTCCCGCTCA
Gibbon                 GCACACACGCATTCCCGCTCA
Ancestral sequences 6  GCACACACGCATTCCCGCTCA
Vervet-AGM             GCACACATGCATTCCCGCTCA
Ancestral sequences 7  GCACACACGCATTCCCGCTCA
Crab-eating macaque    GCACACACGCATTCCCGCTCA
Ancestral sequences 8  GCACACACGCATTCCCGCTCA
Macaque                GCACACACGCATTCCCGCTCA
Ancestral sequences 9  GCACACACGCATTCCCGCTCA
Olive baboon           GCACACACGCATTCCCGCTCA
Ancestral sequences 10 GCACACACGCATTCCCGCTCA
Gelada                 GCACACACGCATTCCCGCTCA
Ancestral sequences 11 GCACACACGCATTCCCGCTCA
Marmoset               GCACACACGCATTCCCGCTTA
Ancestral sequences 12 GCACACACGCATTCCCGCTCA
Mouse Lemur            GCACGCACGCGTTGCCGCCC

rs237897(A/G)

Human                  CCTGCCCACCACTCCTTGCAG
Ancestral sequences 1  CCTGCCCACCGCTCCTTGCAG
Bonobo                 CCTGCCCACCACTCCTTGCAG
Ancestral sequences 2  CCTGCCCACCGCTCCTTGCAG
Chimpanzee             CCTGCCCACCGCTCCTTGCAG
Ancestral sequences 3  CCTGCCCACCGCTCCTTGCAG
Gorilla                CCTGCCCACCGCTCCTTGCAG
Ancestral sequences 4  CCTGCCCACCGCTCCTTGCAG
Orangutan              CCTGCCCACCGCTCCTTGCAG
Ancestral sequences 5  CCTGCCCACCGCTCCTTGCAG
Gibbon                 ACTGCCCACCGCTCCTTGCAG
Ancestral sequences 6  CCTGCCCACCGCTCCTTGCAG
Vervet-AGM             CCTGCCCACCGCTTCTTGCAG
Ancestral sequences 7  CCTGCCCACCGCTTCTTGCAG
Crab-eating macaque    CCTGCCCACCGCTTTTTGCAG
Ancestral sequences 8  CCTGCCCACCGCTTTTTGCAG
Macaque                CCTGCCCACCGCTTTTTGCAG
Ancestral sequences 9  CCTGCCCACCGCTTCTTGCAG
Olive baboon           CCTGCCCACCAGTTCTTGCAG
Ancestral sequences 10 CCTGCCCACCACTTCTTGCAG
Gelada                 CCTGCCCACCACTTCTTGCAG
Ancestral sequences 11 CCTGCCCACCGCTCCTTGCAG
Marmoset               CCTACCCAGCGCTCCCTGCAG
Ancestral sequences 12 CCTGCCCACCGCTCCT--CAG
Mouse Lemur            CTAGCCCTCAGCCCCA--CCG


rs11131149(A/G)

Human                  AAAAAATCGTGCTCTAAACCA
Ancestral sequences 1  AAAAAATCGTGCTCTAAACCA
Bonobo                 AAAAAATCGTGCTCTAAACCA
Ancestral sequences 2  AAAAAATCGTGCTCTAAACCA
Chimpanzee             AAAAAATCGTGCTCTAAACCA
Ancestral sequences 3  AAAAAATCGTGCTCTAAACCA
Gorilla                AAAAAATCGTGCTGCAAACCA
Ancestral sequences 4  AAA-AATCATGCTCTAAACCA
Orangutan              AAA-AATCATGCTCTAAACCA
Ancestral sequences 5  AAA-AATCATGCTCTAAACCA
Gibbon                 AAA-AATCATGCTCTAAACCA
Ancestral sequences 6  AAA-AATCATGCTCTAAACCA
Vervet-AGM             AAA-AATCATACTCTAAACCA
Ancestral sequences 7  AAA-AATCATACTCTAAACCA
Crab-eating macaque    AAA-AATCATACTCTAAACCA
Ancestral sequences 8  AAA-AATCATACTCTAAACCA
                        R
Macaque                AAA-AATCATACTCTAAACCA
Ancestral sequences 9  AAA-AATCATACTCTAAACCA
Olive baboon           AAA-AATCATACTCTAAACCA
Ancestral sequences 10 AAA-AATCATACTCTAAACCA
Gelada                 AAA-AATCATACTCTAAACCA


rs59190448(G/A)

Human                  TGCCATCAGCGGATGTGTACC
Ancestral sequences 1  TGCCATCAGCAGATGTGTACC
Bonobo                 TGCCATCAGCAGATGTGTACC
Ancestral sequences 2  TGCCATCAGCAGATGTGTACC
Chimpanzee             TGCCATCAGCAGATGTGTACC
Ancestral sequences 3  TGCCATCAGCAGATGTGTACC
Gorilla                TGCCATCAGCAGATGTGTACC
Ancestral sequences 4  TGCCATCAGCAGATGTGTACC
Orangutan              TGCCATCAGCAGATGTGTACC
Ancestral sequences 5  TGCCATCAGCAGATGTGTACC
Gibbon                 TGCCATCAGCAGATGTGTACC
Ancestral sequences 6  TGCCATCAGCAGATGTGTACC
Vervet-AGM             TGCCATCAGCAGATGTGTACC
Ancestral sequences 7  TGCCATCAGCAGATGTGTACC
Crab-eating macaque    TGCCATCAGCAGATGTGTACC
Ancestral sequences 8  TGCCATCAGCAGATGTGTACC
Macaque                TGCCATCAGCAGATGTGTACC
Ancestral sequences 9  TGCCATCAGCAGATGTGTACC
Olive baboon           TGCCATCAGCAGATGTGTACC
Ancestral sequences 10 TGCCATCAGCAGATGTGTACC
Gelada                 TGCCATCAGCAGATGTGTACC


rs13316193(C/T)

Human                  ACGGGAATGCTGTTAAATATC
Ancestral sequences 1  ACGGGAATGCTGCTAAATATC
Bonobo                 ACGGGAATGCTGCTAAATATC
Ancestral sequences 2  ACGGGAATGCTGCTAAATATC
Chimpanzee             AGGGGAATGCTGCTAAATATC
Ancestral sequences 3  ACGGGAATGCTGCTAAATATC
Gorilla                ATGGGAATGCTGCTAAATATC
Ancestral sequences 4  ACAGGAATGCTGCTAAATATC
Orangutan              ACAGGAATGCTGCTAAATATC
Ancestral sequences 5  ACAGGAATGCTGCTAAATATC
Gibbon                 ACAGGAATGCTGCTGAATATC
Ancestral sequences 6  ACAGGAATGCTGCTAAATATC
Vervet-AGM             ACAGGAATGCTGCTAAGTATC
Ancestral sequences 7  ACAGGAATGCTGCTAAATATC
Crab-eating macaque    ACAGGAATGCTGCTAAATATC
Ancestral sequences 8  ACAGGAATGCTGCTAAATATC
Macaque                ACAGGAATGCTGCTAAATATC
Ancestral sequences 9  ACAGGAATGCTGCTAAATATC
Olive baboon           ACAGGAATGCTGCTAAATATC
Ancestral sequences 10 ACAGGAATGCTGCTAAATATC
Gelada                 ACAGGAATGCTGCTAAATATC
Ancestral sequences 11 ACAGGAATGCTGCTAAATATC
Marmoset               ACAGGAATGCTGCTAAATATC


rs9872310(G/A)

Human                  CCGTAAGTATAAGTGTTCATA
Ancestral sequences 1  CCGTAAGTATAAGTGTTCATA
Bonobo                 CCGTAAGTATAAGTGTTCACA
Ancestral sequences 2  CCGTAAGTATAAGTGTTCATA
Chimpanzee             CCGTAAGTATAAGTGTTCATA
Ancestral sequences 3  CCGTAAGTATAAGTGTTCATA
Gorilla                CCGTAAGTATAAGTGTTCATA
Ancestral sequences 4  CCATAAGTATAAGTGTTCATA
Orangutan              CCATAAGTATAAGTGTTCATA
Ancestral sequences 5  CCATAAGTATAAGTGTTCATA
Gibbon                 CCATAAGTATAAGTGTTCATG
Ancestral sequences 6  CCATAAGTATAAGTGTTCATA
Vervet-AGM             CCATAAGTATAAGTGTTCATA
Ancestral sequences 7  CCATAAGTATAAGTGTTCATA
Crab-eating macaque    CCATAAGTATAAGTGTTCATA
Ancestral sequences 8  CCATAAGTATAAGTGTTCATA
Macaque                CCATAAGTATAAGTGTTCATA
Ancestral sequences 9  CCATAAGTATAAGTGTTCATA
Olive baboon           CCATAAGTATAAGTGTTCATA
Ancestral sequences 10 CCATAAGTATAAGTGTTCATA
Gelada                 CCATAAGTATAAGTGTTCATA
Ancestral sequences 11 CCATAAGTATAAGTGTTCATA
Marmoset               CCATAAGTACAAGTGTTCATA
Ancestral sequences 12 CCATAAGTATAAGTGTTCATA
Mouse Lemur            CCATAAATGTAGGTGTTCGTA

rs4686302(T/C)

Human                  CCGTAGCAGGCAGCGAGCACG
Ancestral sequences 1  CCGTAGCAGGCAGCGAGCACG
Bonobo                 CCGTAGCAGGCAGCGAGCACG
Ancestral sequences 2  CCGTAGCAGGCAGCGAGCACG
Chimpanzee             CCGTAGCAGGCAGCGAGCACG
Ancestral sequences 3  CCGTAGCAGGCAGCGAGCACG
Gorilla                CCGTAGCAGGCAGCGAGCACG
Ancestral sequences 4  CCGTAGCAGGCAGCGAGCACG
Orangutan              CCGTAGCAGGCAGCGAGCACG
Ancestral sequences 5  CCGTAGCAGGCAGCGAGCACG
Gibbon                 CCGTAGCAGGCAGCGAGCACG
Ancestral sequences 6  CCGTAGCAGGCAGCGAGCACG
Vervet-AGM             CCATAGCAGGCAGCGAGTACG
Ancestral sequences 7  CCATAGCAGGCAGCGAGCACG
Crab-eating macaque    CCATAGCAGGCAGCGAGCACG
Ancestral sequences 8  CCATAGCAGGCAGCGAGCACG
Macaque                CCATAGCAGGCAGCGAGCACG
Ancestral sequences 9  CCATAGCAGGCAGCGAGCACG
Olive baboon           CCATAGCAGGCAGCGAGCACG
Ancestral sequences 10 CCATAGCAGGCAGCGAGCACG
Gelada                 CCATAGCAGGCAGCGAGCACG
Ancestral sequences 11 CCGTAGCAGGCGGCTAGCACG
Marmoset               CCGTAGCAGGCGGCTAGCATG
Ancestral sequences 12 CCGTAGCAGGCGGCCAGCACG
Mouse Lemur            CCGTAGCAGGCGGCCAGCACG


rs237888(T/C)

Human                  CCTAGTTGGATACAGTTATTT
Ancestral sequences 1  CCTAGTTGGACACAGTTATTT
Bonobo                 CCTAGTTGGACACAGTTATTT
Ancestral sequences 2  CCTAGTTGGACACAGTTATTT
Chimpanzee             CCTAGTTGGACACAGTTATTT
Ancestral sequences 3  CCTAGTTG-------------
Gorilla                CCTAGTTG-------------
Ancestral sequences 4  CCTAGTTG-------------
Orangutan              CCTAGTTG-------------
Ancestral sequences 5  CCTAGTTG-------------
Gibbon                 CCTAGTTG-------------
Ancestral sequences 6  CCTAGTTG-------------
Vervet-AGM             CCTAGTTG-------------
Ancestral sequences 7  CCTAGTTG-------------
Crab-eating macaque    CCTAGTTG-------------
Ancestral sequences 8  CCTAGTTG-------------
Macaque                CCTAGTTG-------------
Ancestral sequences 9  CCTAGTTG-------------
Olive baboon           CCTAGTTG-------------
Ancestral sequences 10 CCTAGTTG-------------
Gelada                 CCTAGTTG-------------


rs60902022(C/T)

Human                  CTTCAAAGGATGGGAATCCGT
Ancestral sequences 1  CTTCAAAGGATGGGAATCCGT
Bonobo                 CTTCAAAGGATGGGAATCCGT
Ancestral sequences 2  CTTCAAAGGATGGGAATCCGT
Chimpanzee             CTTCAAAGGATGGGAATCCGT
Ancestral sequences 3  CTTCAAAGGATGGGAATCCGT
Gorilla                CTTCAAAGGATGGGAATCCGT
Ancestral sequences 4  CTTCAAAGGATGGGAATCCAT
Orangutan              CTTCAAAGGATGGGAATCCAT
Ancestral sequences 5  CTTCAAAGGATGGGAATCCAT
Gibbon                 CTTCAAAGGATGGGAATCCAT
Ancestral sequences 6  CTTCAAAGGATGGGAATCCAT
Vervet-AGM             CTTCAAAGGATGGGAATCCAT
Ancestral sequences 7  CTTCAAAGGATGGGAATCCAT
Crab-eating macaque    CTTCAAAGGATGGGAATCCAT
Ancestral sequences 8  CTTCAAAGGATGGGAATCCAT
Macaque                CTTCAAAGGATGGGAATCCAT
Ancestral sequences 9  CTTCAAAGGATGGGAATCCAT
Olive baboon           CTTCAAAGGATGGGAATCCAC
Ancestral sequences 10 CTTCAAAGGATGGGAATCCAT
Gelada                 CTTCAAAGGATGGGAATCCAT
Ancestral sequences 11 CTTCAAAGGATGGGAATCCAT
Marmoset               CTTCAAGGGATGGGAATCCAT


rs6770632(C/A)

Human                  AATTTCTTTCCAATTTT-------------GTAG
Ancestral sequences 1  AATTTCTTTCAAATTTT-------------GTAG
Bonobo                 AATTTCTTTCAAATTTT-------------GTAG
Ancestral sequences 2  AATTTCTTTCAAATTTT-------------GTAG
Chimpanzee             AATTTCTTTCAAATTTT-------------GTAG
Ancestral sequences 3  AATTTCTTTCAAATTTT-------------GTAG
Gorilla                AATTTCTTTCAAATTTT-------------GTAG
Ancestral sequences 4  AATTTCTTTCAAATTTT-------------GTAG
Orangutan              AATTTCTTTCAAATTTT-------------GTAG
Ancestral sequences 5  AATTTCTT----------------------GTAG
Gibbon                 AATTTCTT----------------------GTAG
Ancestral sequences 6  AATTTCTT---------CCAAA----TTTTGTAG
Vervet-AGM             AATTTCTT---------CCAAA----TTTTGTAG
Ancestral sequences 7  AATTTCTT---------CCAAA----TTTTGTAG
Crab-eating macaque    AATTTCTT---------CCAAA----TTTTGTAG
Ancestral sequences 8  AATTTCTT---------CCAAA----TTTTGTAG
Macaque                AATTTCTT---------CCAAA----TTTTGTAG
Ancestral sequences 9  AATTTCTT---------CCAAA----TTTTGTAG
Olive baboon           AATTTCTT---------CCAAA----TTTTGTAG
Ancestral sequences 10 AATTTCTT---------CCAAA----TTTTGTAG
Gelada                 AATTTCTT---------CCAAA----TTTTGTAG
Ancestral sequences 11 AATTTCTT---------TCAAA----TTTTGTAG
Marmoset               AATTTCTT---------TCAAA----TTTTGTAG
Ancestral sequences 12 AATTTCTT---------TCAAA----TTTTGTAG
Mouse Lemur            AATTTCTT---------TCAATAGTAGGTTGTAG


rs237885(T/G)

Human                  CATCTTGTGGTTTAGGTAGGC
Ancestral sequences 1  CATCTTGTGGGTTAGGTAGGC
Bonobo                 CATCTTGTGGGTTAGGTAGGC
Ancestral sequences 2  CATCTTGTGGGTTAGGTAGGC
Chimpanzee             CATCTTGTGGGTTAGGTAGGC
Ancestral sequences 3  CATCTTGTGGGTTAGGTAGGC
Gorilla                CATCTTGTGGGTTAGGTAGGC
Ancestral sequences 4  CATCTTGTGGGTTAGGTAGGC
Orangutan              CATCTTACGGGTTAGGTAGGC
Ancestral sequences 5  CATCTTGTGGGTTAGGTAGGC
Gibbon                 CATCTTGTGGGTTAGGTAGGC
Ancestral sequences 6  CATCTTGTGGGTTAGGTAGGC
Vervet-AGM             CATCTTGTGAGTTACGTAGGC
Ancestral sequences 7  CATCTTGTGAGTTAGGTAGGC
Crab-eating macaque    CATCTTGTGGGTTAGGTAGGC
Ancestral sequences 8  CATCTTGTGAGTTAGGTAGGC
Macaque                CATCTTGTGAGTTAGGTAGGC
Ancestral sequences 9  CATCTTGTGAGTTAGGTAGGC
Olive baboon           CATCTTGTGAGTTAGGTAGTC
Ancestral sequences 10 CATCTTGTGAGTTAGGTAGTC
Gelada                 CATCTTGTGAGTTAGGTAGTC
Ancestral sequences 11 CATCTTGTGGGTTAGGTAGGC
Marmoset               CATCTTGTGGGTTAGGTAGGC
Ancestral sequences 12 ---------GGTTAGGTAGGC
Mouse Lemur            ---------GGCTGGGCGGGG


rs1042778(G/T)
Human                  CCCCAAGGAGGGGAGGGATAC
Ancestral sequences 1  CCCCAAGGAGTGGAGGGATAC
Bonobo                 CCCCAAGGAGTGGAGGGATAC
Ancestral sequences 2  CCCCAAGGAGTGGAGGGATAC
Chimpanzee             CCCCAAGGAGTGGAGGGATAC
Ancestral sequences 3  CCCCAAGGAGTGGAGGGATAC
Gorilla                CCCCAAGTTGTGGAGGGATAT
Ancestral sequences 4  CCCCAAGGAGTGGAGGGATAC
Orangutan              CCCCAAGGAGTGGAGGGATAC
Ancestral sequences 5  CCCCAAGGAGTGGAGGGATAC
Gibbon                 CCCCAAGGAGTGGAGGGATAC
Ancestral sequences 6  CCCCAAGGAGTGGAGGGATAC
Vervet-AGM             CCCCAAGGAGTGGAGGGAAAC
Ancestral sequences 7  CCCCAAGGAGTGGAGGGATAC
Crab-eating macaque    CCCCAAGGAGTGGAGGGATAC
Ancestral sequences 8  CCCCAAGGAGTGGAGGGATAC
Macaque                CCCCAAGGAGTAGAGGGATAC
Ancestral sequences 9  CCCCAAGGAGTGGAGGGATAC
Olive baboon           CCCCAAGGAGTGGAGGGATAC
Ancestral sequences 10 CCCCAAGGAGTGGAGGGATAC
Gelada                 CCCCAAGGAGTGGAGGGATAC
Ancestral sequences 11 CCCCAAGGAGTGGAGGGATAC
Marmoset               CCCCAAGGAGTGGAGCGATAC
Ancestral sequences 12 CCCCAAGGAGTGGAGGGATAC
Mouse Lemur            CCCCCAGGAGCGGAGGCATTC

rs237911(G/A)

Human                  GGATCTGCTGGGTCC-ACCCTG
Ancestral sequences 1  GGATCTGCTGAGTCC-ACCCTG
Bonobo                 GGATCTGCTGAGTCC-ACCCTG
Ancestral sequences 2  GGATCTGCTGAGTCC-ACCCTG
Chimpanzee             GGATCTGCTGAGTCC-ACCCTG
Ancestral sequences 3  GGATCTGCTGAGTCC-ACCCTG
Gorilla                GGATCTGCTGAGTCC-ACCCTG
Ancestral sequences 4  GGATCTGCTGGGTCC-ACCCTG
Orangutan              GGATCTGCTGGGTCC-ACCCTG
Ancestral sequences 5  GGATCTGCTGGGTCC-ACCCTG
Gibbon                 GGATCTGCTGGGTCC-ACCCTG
Ancestral sequences 6  GGATCTGCTGGGTCC-ACCCTG
Vervet-AGM             GGATCTGCTGGGTCC-ACCCTG
Ancestral sequences 7  GGATCTGCTGGGTCC-ACCCTG
Crab-eating macaque    GGATCTGCTGGGTCC-ACCCTG
Ancestral sequences 8  GGATCTGCTGGGTCC-ACCCTG
Macaque                GGATCTGCTGGGTCC-ACCCTG
Ancestral sequences 9  GGATCTGCTGGGTCC-ACCCTG
Olive baboon           GGATCTGCTGGGTCC-ACCCTG
Ancestral sequences 10 GGATCTGCTGGGTCC-ACCCTG
Gelada                 GGATCTGCTGGGTCC-ACCCTG
Ancestral sequences 11 GGATCTGCTGGGTCC-ACCCTG
Marmoset               AGATCTGCTGGGTCT-ACCCTG
Ancestral sequences 12 GGATCTGCTGGGTCC-ACCCTG
Mouse Lemur            GGCTCTGCCGGGTCCCAGCCTG


rs2254298(A/G)

Human                  CCG-CAAACTGGGAAAACAG--GG
Ancestral sequences 1  CCG-CAAACTGGGAAAACAG--GG
Bonobo                 CCG-CAAACTGGGAAAACAG--GG
Ancestral sequences 2  CCG-CAAACTGGGAAAACAG--GG
Chimpanzee             CCG-CAAACTGGGAAAACAG--GG
Ancestral sequences 3  CCG-CAAACTGGGAAAACAG--GG
Gorilla                CCA-CAAACTGGGAAAACAG--GG
Ancestral sequences 4  CCG-CAAACTGGGAAAACAG--GG
Orangutan              CCG-CAAACTGGGAAAACAG--GG
Ancestral sequences 5  CCG-CAAACTGGGAAAACAG--GG
Gibbon                 CCG-CAAACTGGGAAAACAG--GG
Ancestral sequences 6  CCTCCAAACTGGGAAAACAG----
Vervet-AGM             CCTCCAAAGTGGGAAAACAG----
Ancestral sequences 7  CCTCCAAAGTGGGAAAACAG----
Crab-eating macaque    CCTCCAAAGTGGGAAAACAGGG--
Ancestral sequences 8  CCTCCAAAGTGGGAAAACAGGG--
                              R
Macaque                CCTCCAAAGTGGGAAAACAGGG--
Ancestral sequences 9  CCTCCAAAGTGGGAAAACAGGG--
Olive baboon           CCTCCAAAGTGGGAAAACAGGG--
Ancestral sequences 10 CCTCCAAAGTGGGAAAACAGGG--
Gelada                 CCTCCAAAGTGGGAAAACAGGG--
Ancestral sequences 11 CCTCCAAACTGGGAAAACAG----
Marmoset               CCCCTAAACTGGAAAATCAG----

rs53576(A/G)
Human                  ATGCCCGAGGATCCTCAGTCC
Ancestral sequences 1  ATGCCCGAGGGTCCTCAGTCC
Bonobo                 ATGCCCGAGGGTCCTCAGTCC
Ancestral sequences 2  ATGCCCGAGGGTCCTCAGTCC
Chimpanzee             ATGCCCGAGGGTCCTCAGTCC
Ancestral sequences 3  ATGCCCGAGGGTCCTCAGTCC
Gorilla                ATGCCCGAGGGTCCTCAGTCC
Ancestral sequences 4  ATGCCCGAGGGTCCTCAGTCC
Orangutan              ATGCCCGAGGGTCCTCAGTCC
Ancestral sequences 5  ATGCCAGAGGGTCCTCAGTCC
Gibbon                 ATGCCAGAGGGTCCTCAGTCC
Ancestral sequences 6  ATGCCAGAGGGTCCTCAGTCC
Vervet-AGM             ATGCCAGATGGTCCTCAGTCC
Ancestral sequences 7  ATGCCAGATGGTCCTCAGTCC
Crab-eating macaque    ATGCCAGATGGTCCTCAGTCC
Ancestral sequences 8  ATGCCAGATGGTCCTCAGTCC
Macaque                ATGCCAGATGGTCCTCAGTCC
Ancestral sequences 9  ATGCCAGATGGTCCTCAGTCC
Olive baboon           ATGCCAGATGGTCCTCAGTCC
Ancestral sequences 10 ATGCCAGATGGTCCTCAGTCC
Gelada                 ATGCCAGATGGTCCTCAGTCC
Ancestral sequences 11 ATGCCAGAGGGTCCTCAGTCC
Marmoset               GTGCCAGAGGATCCTCAGTCC


rs2268490(T/C)

Human                  CACTG--TTTTGCCTAGTTGGAT
Ancestral sequences 1  CACTG--TTTTGCCTAGTTGGAC
Bonobo                 CACTG--TTTTGCCTAGTTGGAC
Ancestral sequences 2  CACTG--TTTTGCCTAGTTGGAC
Chimpanzee             CACTG--TTTTGCCTAGTTGGAC
Ancestral sequences 3  CACTGTGTTTTGCCTAGTTG---
Gorilla                CACTGTGTTTTGCCTAGTTG---
Ancestral sequences 4  CACTGTGTTTTGCCTAGTTG---
Orangutan              CACTGTGTTTTGCCTAGTTG---
Ancestral sequences 5  CACTGTGTTTTGCCTAGTTG---
Gibbon                 CACTGAGTTTTGCCTAGTTG---
Ancestral sequences 6  CACTGTGTTTTGCCTAGTTG---
Vervet-AGM             CACCGTGTTTTGCCTAGTTG---
Ancestral sequences 7  CACCGTGTTTTGCCTAGTTG---
Crab-eating macaque    CACCGTGTTTTGCCTAGTTG---
Ancestral sequences 8  CACCGTGTTTTGCCTAGTTG---
Macaque                CACCGTGTTTTGCCTAGTTG---
Ancestral sequences 9  CACCGTGTTTTGCCTAGTTG---
Olive baboon           CACCGTGTTTTGCCTAGTTG---
Ancestral sequences 10 CACCGTGTTTTGCCTAGTTG---
Gelada                 CACCGTGTTTTGCCTAGTTG---


rs2268493(C/T)

Human                  AAGAAATGAATAAAGTAACTG
Ancestral sequences 1  AAGAAATGAATAAAGTAACTG
Bonobo                 AAGAAATGAATAATGTCACTG
Ancestral sequences 2  AAGAAATGAATAATGTCACTG
Chimpanzee             AAGAAATGAATAATGTCACTG
Ancestral sequences 3  AAGAAATGAATAAAGTAACTG
Gorilla                AAGAAATGAATAAAGTAACTG
Ancestral sequences 4  AAGAAATGAATAAAGTAACTG
Orangutan              AAGAAATGAATAAAGTAACTG
Ancestral sequences 5  AAGAAATGAATAAAGTAACTG
Gibbon                 AAGAAATGAATGAAGTAACTG
Ancestral sequences 6  AAGAAATGAATAAAGTAACTG
Vervet-AGM             AAGAAATGAATAAAGTAACCG
Ancestral sequences 7  AAGAAATGAATAAAGTAACCG
Crab-eating macaque    AAGAAATGAATAAAGTAACCA
Ancestral sequences 8  AAGAAATGAATAAAGTAACCG
Macaque                AAGAAATGAATAAAGTAACCG
Ancestral sequences 9  AAGAAATGAATAAAGTAACCG
Olive baboon           AAGAAATGAATAAAGTAACCA
Ancestral sequences 10 AAGAAATGAATAAAGTAACCG
Gelada                 AAGAAATGAATAAAGTAACCG
Ancestral sequences 11 AAGAAATGAATAAAGTAACTG
Marmoset               AAGAAATGAATAAAGTAACTG


rs237917(T/C)

Human                  CTAACTTAGTTTTAATCTAAA
Ancestral sequences 1  CTAATTTAGTCTTAATCTAAA
Bonobo                 CTAATTTAGTCTTAATCTAAA
Ancestral sequences 2  CTAATTTAGTCTTAATCTAAA
Chimpanzee             CTAATTTAGTCTTAATCTAAA
Ancestral sequences 3  CTAATTTAGTCTTAATCTAAA
Gorilla                CTAATTTAGTCTTAATCTAAA
Ancestral sequences 4  CTAATTTAGTCTTAATCTAAA
Orangutan              CTAATTTAGTCTTAATCTAAA
Ancestral sequences 5  CTAATTTAGTCTTAATCTAAA
Gibbon                 CTAATTTAGTCTTAATCTAAA
Ancestral sequences 6  CTAATTTAGTCTTAATCTAAA
Vervet-AGM             CTAATTTAGTCTTAATCTAAA
Ancestral sequences 7  CTAATTTAGTCTTAATCTAAA
Crab-eating macaque    CTAATTTAGTCTTAATCTAAA
Ancestral sequences 8  CTAATTTAGTCTTAATCTAAA
Macaque                CTAATTTAGTCTTAATCTAAA
Ancestral sequences 9  CTAATTTAGTCTTAATCTAAA
Olive baboon           CTAATTTAGTCTTAATCTAAA
Ancestral sequences 10 CTAATTTAGTCTTAATCTAAA
Gelada                 CTAATTTAGTCTTAATCTAAA
Ancestral sequences 11 CTAATTTAGCCTTAATCTTAA
Marmoset               CTAATTCAGCCTTGATCTTAA
Ancestral sequences 12 CTAATTTAGCCTTAATCTTAA
Mouse Lemur            CTAATTTGGCCTTCCTCTTAA


rs237889(T/C)
Human                  AGCAAGGCCATAGGAACTTGT
Ancestral sequences 1  AGCAAGGCCACAGGAACTTGT
Bonobo                 AGCAAGGCCACAGGAACTTGT
Ancestral sequences 2  AGCAAGGCCACAGGAACTTGT
Chimpanzee             AGCAAGGCCACAGGAACTTGT
Ancestral sequences 3  AGCAAGGCCACAGGAACTTGT
Gorilla                AGCAAGGCCACAGGAACTTGT
Ancestral sequences 4  AGCAAGGCCACAGGAACTTGT
Orangutan              AGCAAGGCCACAGGAACTTGT
Ancestral sequences 5  AGCAAGGCCACAGGAACTTGT
Gibbon                 GGCAAGGCCACAGGAACTTGT
Ancestral sequences 6  AGCAAGGCCACAGGAACTTGT
Vervet-AGM             AGCAAGGCCACAGGAACTTGT
Ancestral sequences 7  AGCAAGGCCACAGGAACTTGT
Crab-eating macaque    AGCAAGGCCACAGGAACTTGT
Ancestral sequences 8  AGCAAGGCCACAGGAACTTGT
Macaque                AGCAAGGCCACAGGAACTTGT
Ancestral sequences 9  AGCAAGGCCACAGGAACTTGT
Olive baboon           AGCAAGGCCACAGGAACTTGT
Ancestral sequences 10 AGCAAGGCCACAGGAACTTGT
Gelada                 AGCAAGGCCACAGGAACTTGT
Ancestral sequences 11 AGCAAGGCCACAGGAACTTGT
Marmoset               AGCAAAGCCACAGGAGCTGGT


VTR1A

rs1042615(A/G)
Human                  AGGCCGACGCA-AACATGCCGA
Ancestral sequences 1  AGGCCGACGCG-AACATGCCGA
Bonobo                 AGGCCGACGCG-AACATGCCGA
Ancestral sequences 2  AGGCCGACGCG-AACATGCCGA
Chimpanzee             AGGCCGACGCG-AACATGCCGA
Ancestral sequences 3  AGGCCGACGCG-AACATGCCGA
Gorilla                AGGCCGACGCG-AACATGCCGA
Ancestral sequences 4  AGGCCGACGCG-AACATGCCGA
Orangutan              AGGCCGACGCG-AACATGCCGA
Ancestral sequences 5  AGGCCGACGCG-AACATGCCGA
Gibbon                 AGGCCGACGCGAAACATGCCGA
Ancestral sequences 6  AGGCCGACGCG-AACATGCCGA
Vervet-AGM             AGGCCGACGCG-AACATGCCGA
Ancestral sequences 7  AGGCCGACGCG-AACATGCCGA
Crab-eating macaque    AGGCCGACGCG-AACATGCCGA
Ancestral sequences 8  AGGCCGACGCG-AACATGCCGA
Macaque                AGGCCGACGCG-AACATGCCGA
Ancestral sequences 9  AGGCCGACGCG-AACATGCCGA
Olive baboon           AGGCCGACGCG-AACATGCCGA
Ancestral sequences 10 AGGCCGACGCG-AACATGCCGA
Gelada                 AGGCCGACGCG-AACATGCCGA
Ancestral sequences 11 AGGCCGACGCG-AACATGCCGA
Marmoset               AGGCCGATGCG-AACATGCCAA
Ancestral sequences 12 AGGCCGACGCG-AACATGCCGA
Mouse Lemur            AGGCCGACGCG-AACATGCCGA

rs3803107(A/G)

Human                  GAAAATAAAAG-----AAACTAACAA
Ancestral sequences 1  GAAAATAAAAG-----AAACCAACAA
Bonobo                 GAAAATAAAAG-----AAACCAACAA
Ancestral sequences 2  GAAAATAAAAG-----AAACCAACAA
Chimpanzee             GAAAATAAAAG-----AAACCAACAA
Ancestral sequences 3  GAAAATAAAAG-----AAACCAACAA
Gorilla                GAAAATAAAAG-----AAACCAACAA
Ancestral sequences 4  GAAAATAAAAG-----AAACCAACAA
Orangutan              GAAAATAAAAG-----AAACCAACAA
Ancestral sequences 5  GAAAATAAAAG-----AAACCAACAA
Gibbon                 GAAAATAAAAG-----AAACCAACAA
Ancestral sequences 6  GAAAATAAAAG-----AAACCAACAA
Vervet-AGM             GAAAATAAAAG-----AAACCAACAA
Ancestral sequences 7  GAAAATAAAAG-----AAACCAACAA
Crab-eating macaque    GAAAAT----------AAACCAACAA
Ancestral sequences 8  GAAAAT----------AAACCAACAA
Macaque                GAAAAT----------AAACCAACAA
Ancestral sequences 9  GAAAAT----------AAACCAACAA
Olive baboon           GAAAAT-----AAAAGAAACCAACAA
Ancestral sequences 10 GAAAAT-----AAAAGAAACCAACAA
Gelada                 GAAAAT-----AAAAGAAACCAACAA
Ancestral sequences 11 GAAAATAAAAG-----AAACCAACAA
Marmoset               GAAAATAAAAG-----AAACCAATAA
Ancestral sequences 12 GAAAATAAAAG-----AAACCAACAA
Mouse Lemur            AAAATTAAAAG-----AAATCAACAA


rs10784339(C/G)

Human                  A-AATACAACTGGGTAGGGTGA
Ancestral sequences 1  A-AATACAACTGGGTAGGGTGA
Bonobo                 A-AATACGACTGGGTAGGGTGA
Ancestral sequences 2  A-AATACAACTGGGTAGGGTGA
Chimpanzee             A-AATACAACTGGGTAGGGTGA
Ancestral sequences 3  A-AATACAACTGGGTAGGGTGA
Gorilla                A-AATACAACTGGGTAGGGTGA
Ancestral sequences 4  A-AATACAACTGGGTAGGGTGA
Orangutan              A-AATACAACTGGGTAGAGTGA
Ancestral sequences 5  A-AATACAACTGGGTAGGGTGA
Gibbon                 A-AATACAGCAGGGTAGGGTGA
Ancestral sequences 6  A-AATACAACTGGGTAGGGTGA
Vervet-AGM             A-AAAACAACTGGGTAGGGTGA
Ancestral sequences 7  A-AAAACAACTGGGTAGGGTGA
Crab-eating macaque    A-AAAACAACTGGGTAGGGTGA
Ancestral sequences 8  A-AAAACAACTGGGTAGGGTGA
Macaque                AAAAAACAACTGGGTAGGGTGA
Ancestral sequences 9  A-AAAACAACTGGGTAGGGTGA
Olive baboon           A-AAAACAACTGGGTAGGGTGA
Ancestral sequences 10 A-AAAACAACTGGGTAGGGTGA
Gelada                 A-AAAACAACTGGGTAGGGTGA
Ancestral sequences 11 A-AATACAACTGGGTAGGGTGA
Marmoset               A-AATACAGCTGGGTAGATTGA

rs11174811(A/C)

Human                  TCATGCTTTTCTTGACCAATA
Ancestral sequences 1  TCATGCTTTTCTTGACCAATA
Bonobo                 TCATGCTTTTCTTGACCAATA
Ancestral sequences 2  TCATGCTTTTCTTGACCAATA
Chimpanzee             TCATGCTTTTCTTGACCAATA
Ancestral sequences 3  TCATGCTTTTCTTGACCAATA
Gorilla                TCATGCTTTTCTTGACCAATA
Ancestral sequences 4  TCATGCTTTTCTTGACCAATA
Orangutan              TCATGCTTTTCTTGACCAATA
Ancestral sequences 5  TCATGCTTTTCTTGACCAATA
Gibbon                 TCATACTTTTCTTGACCAATA
Ancestral sequences 6  TCATGCTTTTCTTGACCAATA
Vervet-AGM             TCATGCTTTTCTTTACCAGTA
Ancestral sequences 7  TCATGCTTTTCTTTACCAGTA
Crab-eating macaque    TCATGCTTTTCTTTACCAGTA
Ancestral sequences 8  TCATGCTTTTCTTTACCAGTA
Macaque                TCATGCTTTTCTTTACCAGTA
Ancestral sequences 9  TCATGCTTTTCTTTACCAGTA
Olive baboon           TCATGCTTTTCTTTACCAGTA
Ancestral sequences 10 TCATGCTTTTCTTTACCAGTA
Gelada                 TCATGCTTTTCTTTACCAGTA
Ancestral sequences 11 TCATGCTTTTCTTGACCAATA
Marmoset               TTGTGCTTTCCTTGACCAATA
Ancestral sequences 12 TCATGCTTTTCTTGACCAATA
Mouse Lemur            TCATGCTTTGTGTGGCCAATA


rs3021529(A/G)
Human                  TTCCCACAGCGGGGATGGCGG
Ancestral sequences 1  TTCCCACAGCGGGGATGGCGG
Bonobo                 TTCCCACAGCGGGGATGGCGG
Ancestral sequences 2  TTCCCACAGCGGGGATGGCGG
Chimpanzee             TTCCCACAGCGGGGATGGCGG
Ancestral sequences 3  TTCCCACAGCGGGGATGGCGG
Gorilla                TTCCCACAGCGGTGATGGCGG
Ancestral sequences 4  TTCCCACAGCGGGGATGGCGG
```

Orangutan	TTCCCACAGC**G**GGGATGGC[G](https://useast.ensembl.org/Pongo_abelii/ZMenu/TextSequence?db=core;factorytype=Location;r=12:63151400-63152400;vdb=variation;vf=2280975" \o "G/A)G

```
Ancestral sequences 5  TTCCCACAGCGGGGATGGCGG
Gibbon                 TTCCCACAGCGGGGATGGCGG
Ancestral sequences 6  TTCCCACAGCGGGGATGGCGG
Vervet-AGM             TTCCCACAGCGGGGATGGCGG
Ancestral sequences 7  TTCCCACAGCGGGGATGGCGG
Crab-eating macaque    TTCCCACAGCGGGGATGGCGG
Ancestral sequences 8  TTCCCACAGCGGGGATGGCGG
```

Macaque	TTCCCACAGC**G**[G](https://useast.ensembl.org/Macaca_mulatta/ZMenu/TextSequence?db=core;factorytype=Location;r=12:63151400-63152400;vdb=variation;vf=6298353" \o "G/C)GGATGGCGG

```
Ancestral sequences 9  TTCCCACAGCGGGGATGGCGG
Ancestral sequences 10 TTCCCACAGCGGGGATGGCGG
Gelada                 TTCCCACAGCGGGGATGGCGG
Ancestral sequences 11 TTCCCACAGCGGGGATGGCGG
Marmoset               TTCCCATAGCGGGGATAGCGG
Ancestral sequences 12 TTCCCACAGCGGGGATGGCGG
Mouse Lemur            TTCCCACAGCGGAAATGGCGG


rs10877969(T/C)

Human                  CTTT---GTT-TAATCCATATAGTT
Ancestral sequences 1  CTTT---TTT-TAACCCATATAGTT
Bonobo                 CTTT---TTT-TAACCCATAGAGTT
Ancestral sequences 2  CTTT---TTT-TAACCCATAGAGTT
Chimpanzee             CTTT---TTT-TAACCCATAGAGTT
Ancestral sequences 3  CTTT---TTT-TAACCCATATAGTT
Gorilla                CTTT---TTT-TAACCCATATAGTT
Ancestral sequences 4  CTTTTTTTTT-TAACCCATATAGTT
Orangutan              CTTTTTTTTT-TAACCCATATAGTT
Ancestral sequences 5  CTTTTTTTTT-AAACCCATATAGTT
Gibbon                 CTTTTTTTTT-AAACCCATA-TGTT
Ancestral sequences 6  CTTT---TTT-AAACCCATATAGTT
Vervet-AGM             CTTT---TTTTAAAGCCATATAGTC
Ancestral sequences 7  CTTT---TTTAAAAGCCATATAGTT
Crab-eating macaque    CTTT---TTTAAAAGCCATATAGTT
Ancestral sequences 8  CTTT---TTTAAAAGCCATATAGTT
Macaque                CTTT---TTTAAAAGCCATATAGTT
Ancestral sequences 9  CTTT---TTTAAAAGCCATATAGTT
Olive baboon           CTTT---TTTAAAAGCCATATAGTT
Ancestral sequences 10 CTTT---TTTAAAAGCCATATAGTT
Gelada                 CTTT---TTTAAAAGCCATGTAGTT
Ancestral sequences 11 CTTT----TT-AAACCCATATAGTT
Marmoset               CTTT----TT-AAACCCATATAGTT


rs3759292(G/A)
```

Human	ACTATTAC-[A](https://useast.ensembl.org/Homo_sapiens/ZMenu/TextSequence?db=core;factorytype=Location;r=12:63153033-63154033;vdb=variation;vf=646517675" \o "A/G)T**A**-T[G](https://useast.ensembl.org/Homo_sapiens/ZMenu/TextSequence?db=core;factorytype=Location;r=12:63153033-63154033;vdb=variation;vf=492912712" \o "G/A)AGGCA[C](https://useast.ensembl.org/Homo_sapiens/ZMenu/TextSequence?db=core;factorytype=Location;r=12:63153033-63154033;vdb=variation;vf=627217685" \o "C/T)TA

```
Ancestral sequences 1  ACTATTAC-ATA-TGAGGCACTA
Bonobo                 ACTATTAC-ATA-TGAGGCACTA
Ancestral sequences 2  ACTATTAC-ATA-TGAGGCACTA
Chimpanzee             ACTATTAC-ATA-TGAGGCACTA
Ancestral sequences 3  ACTATTAC-ATA-TGAGGCACTA
Gorilla                ACTATTAC-ATA-TGAGGCACTA
Ancestral sequences 4  ACTATTACAATATAGACTCACTA
Orangutan              ACTATTACAATATAGACTCACTA
Ancestral sequences 5  ACTATTACAATATAGACTCACTA
Gibbon                 ACCATTACAATATAGACTCACTA
Ancestral sequences 6  ACTATTACAATATAGACTCACTA
Vervet-AGM             ACTATTACAATATAGACTCACTA
Ancestral sequences 7  ACTATTACAATATAGACTCACTA
Crab-eating macaque    ACTATTACAATATAGATTCACTA
Ancestral sequences 8  ACTATTACAATATAGATTCACTA
```

Macaque	ACTATTACAAT**A**TAGA[T](https://useast.ensembl.org/Macaca_mulatta/ZMenu/TextSequence?db=core;factorytype=Location;r=12:63153033-63154033;vdb=variation;vf=6298394" \o "T/C)TCACTA

```
Ancestral sequences 9  ACTATTACAATATAGACTCACTA
Olive baboon           ACTATTACAATATAGACTCACTA
Ancestral sequences 10 ACTATTACAATATAGACTCACTA
Gelada                 ACTATTACAATATAGACTCACTA
Ancestral sequences 11 ACTATTACAATATAGACTCACTA
Marmoset               ACCATTACAATACAGACTTACTA


VTR1B
rs28676508(T/C)
```

Human	[T](https://useast.ensembl.org/Homo_sapiens/ZMenu/TextSequence?db=core;factorytype=Location;r=1:206109845-206110845;vdb=variation;vf=27045367" \o "T/-)[G](https://useast.ensembl.org/Homo_sapiens/ZMenu/TextSequence?db=core;factorytype=Location;r=1:206109845-206110845;vdb=variation;vf=135736865" \o "G/T)[T](https://useast.ensembl.org/Homo_sapiens/ZMenu/TextSequence?db=core;factorytype=Location;r=1:206109845-206110845;vdb=variation;vf=135736871" \o "T/C)[GG](https://useast.ensembl.org/Homo_sapiens/ZMenu/TextSequence?db=core;factorytype=Location;r=1:206109845-206110845;vdb=variation;vf=26529267" \o "GGCGGCTCGAGAGG/-)[C](https://useast.ensembl.org/Homo_sapiens/ZMenu/TextSequence?db=core;factorytype=Location;r=1:206109845-206110845;vdb=variation;vf=26529267;vf=5107748" \o "GGCGGCTCGAGAGG/- C/A/G/T)[G](https://useast.ensembl.org/Homo_sapiens/ZMenu/TextSequence?db=core;factorytype=Location;r=1:206109845-206110845;vdb=variation;vf=26529267;vf=1372981" \o "GGCGGCTCGAGAGG/- G/A)[G](https://useast.ensembl.org/Homo_sapiens/ZMenu/TextSequence?db=core;factorytype=Location;r=1:206109845-206110845;vdb=variation;vf=26529267" \o "GGCGGCTCGAGAGG/-)[C](https://useast.ensembl.org/Homo_sapiens/ZMenu/TextSequence?db=core;factorytype=Location;r=1:206109845-206110845;vdb=variation;vf=26529267;vf=27110211" \o "GGCGGCTCGAGAGG/- C/T)[T](https://useast.ensembl.org/Homo_sapiens/ZMenu/TextSequence?db=core;factorytype=Location;r=1:206109845-206110845;vdb=variation;vf=26529267" \o "GGCGGCTCGAGAGG/-)**C**[G](https://useast.ensembl.org/Homo_sapiens/ZMenu/TextSequence?db=core;factorytype=Location;r=1:206109845-206110845;vdb=variation;vf=26529267;vf=26993446" \o "GGCGGCTCGAGAGG/- G/A/C)[A](https://useast.ensembl.org/Homo_sapiens/ZMenu/TextSequence?db=core;factorytype=Location;r=1:206109845-206110845;vdb=variation;vf=26529267;vf=26563355" \o "GGCGGCTCGAGAGG/- A/C)[GA](https://useast.ensembl.org/Homo_sapiens/ZMenu/TextSequence?db=core;factorytype=Location;r=1:206109845-206110845;vdb=variation;vf=26529267" \o "GGCGGCTCGAGAGG/-)[G](https://useast.ensembl.org/Homo_sapiens/ZMenu/TextSequence?db=core;factorytype=Location;r=1:206109845-206110845;vdb=variation;vf=26529267;vf=27102519" \o "GGCGGCTCGAGAGG/- G/A)[G](https://useast.ensembl.org/Homo_sapiens/ZMenu/TextSequence?db=core;factorytype=Location;r=1:206109845-206110845;vdb=variation;vf=26529267;vf=27025118" \o "GGCGGCTCGAGAGG/- G/T)[C](https://useast.ensembl.org/Homo_sapiens/ZMenu/TextSequence?db=core;factorytype=Location;r=1:206109845-206110845;vdb=variation;vf=139251924" \o "C/T)[T](https://useast.ensembl.org/Homo_sapiens/ZMenu/TextSequence?db=core;factorytype=Location;r=1:206109845-206110845;vdb=variation;vf=26588701" \o "T/A)G[C](https://useast.ensembl.org/Homo_sapiens/ZMenu/TextSequence?db=core;factorytype=Location;r=1:206109845-206110845;vdb=variation;vf=27221109;vf=27145776" \o "C/A CCGTC/C)

```
Ancestral sequences 1  TGTGGCGGCTCGAGAGGCTGC
Bonobo                 TGTGGCGGCTCGAGAGGCTGC
Ancestral sequences 2  TGTGGCGGCTCGAGAGGCTGC
Chimpanzee             TGTGGCGGCTCGAGAGGCTGC
Ancestral sequences 3  TGTGGCGGCTCGAGAGGCTGC
Gorilla                TGTGGCGGCTCGAGAGGCTGC
Ancestral sequences 4  TGTGGCGGCTCGAGAGGCTGC
Orangutan              TGTGGCGGCTCGAGAGGCTGC
Ancestral sequences 5  TGTGGCGGCTCGAGAGGCTGC
Gibbon                 TGTGGCGGCTCGAGAGGCTGC
Ancestral sequences 6  TGTGGCGGCTCGAGAGGCTGC
Vervet-AGM             TGTGGCGGCTCGAAAGGCTGC
Ancestral sequences 7  TGTGGCGGCTCGAGAGGCTGC
Crab-eating macaque    TGTGGCGGCTCGAGAGGCTGC
Ancestral sequences 8  TGTGGCGGCTCGAGAGGCTGC
Macaque                TGTGGCGGCTCGAGAGGCTGC
Ancestral sequences 9  TGTGGCGGCTCGAGAGGCTGC
Olive baboon           TGTGGCGGCTCGAGAGGCTGC
Ancestral sequences 10 TGTGGCGGCTCGAGAGGCTGC
Gelada                 TGTGGCGGCTCGAGAGGCTGC
Ancestral sequences 11 TGTGGCGGCTCGAGAGGCTGC
Marmoset               TGTGGCGGCTCGAGAGGCTGC
Ancestral sequences 12 TGTGGCGGCTCGAGAGGCTGC
Mouse Lemur            TGCGGCGGCTCGACAGGCTGC

rs28632197(T/C)
Human                  GAGCCGCCGGCGCATCCTGGG
Ancestral sequences 1  GAGCCGCCGGCGCATCCTGGG
Bonobo                 GAGCCGCCGGCGCATCCTGGG
Ancestral sequences 2  GAGCCGCCGGCGCATCCTGGG
Chimpanzee             GAGCCGCCGGCGCATCCTGGG
Ancestral sequences 3  GAGCCGCCGGCGCATCCTGGG
Gorilla                GAGCCGCCGGCGCATCCTGGG
Ancestral sequences 4  GAGCCGCCGGCGCATCCTGGG
Orangutan              GAGCCGCCGGCGCATCTTGGG
Ancestral sequences 5  GAGCCGCCGGCGCATCCTGGG
Gibbon                 GAGCCGCCGGCGCATCCTGGG
Ancestral sequences 6  GAGCCGCCGGCGCATCCTGGG
Vervet-AGM             GAGCCGCCGGCACATCCTGGG
Ancestral sequences 7  GAGCCGCCGGCACATCCTGGG
Crab-eating macaque    GAGCCGCCGGCACATCCTGGG
Ancestral sequences 8  GAGCCGCCGGCACATCCTGGG
Macaque                GAGCCGCCGGCACATCCTGGG
Ancestral sequences 9  GAGCCGCCGGCACATCCTGGG
Olive baboon           GAGCCGCCGGCACATCCTGGG
Ancestral sequences 10 GAGCCGCCGGCACATCCTGGG
Gelada                 GAGCCGCCGGCACATCCTGGG
Ancestral sequences 11 GAGCCGCCGGCGCATCCGGGG
Marmoset               GAGCCGCCGGCGCATCCGGGG
Ancestral sequences 12 GAGCCGCCGGCGCATCCGGGG
Mouse Lemur            GAGCTGCCGGCGCATGCGTGG

rs33985287(T/C)
Human                  CGCTCCGCTTT--AGACAGGGCT
Ancestral sequences 1  CGCTCCGCTTC--AGACAGGGCT
Bonobo                 CGCTCCGCTTC--AGACAGGGCT
Ancestral sequences 2  CGCTCCGCTTC--AGACAGGGCT
Chimpanzee             CGCTCCGCTTC--AGACAGGGCT
Ancestral sequences 3  CACTCCGCTTC--AGACAGGGCT
Gorilla                CACTCCGCTTC--AGACAGGGCT
Ancestral sequences 4  CACTCCGCTTC--AGACAGGGCT
Orangutan              CACTCCGCTTC--AGACAGGGCT
Ancestral sequences 5  CACTCCGCTTC--AGACAGGGCT
Gibbon                 CACTCCGCTTC--AGACAGGGCT
Ancestral sequences 6  CACCCCGCTTC--AGACAGGGCT
Vervet-AGM             CACCCTGCTTC--AGACAGGGCT
Ancestral sequences 7  CACCCCGCTTC--AGACAGGGCT
Crab-eating macaque    CACCCCGCTTC--AGACAGGGCT
Ancestral sequences 8  CACCCCGCTTC--AGACAGGGCT
Macaque                CACCCCGCTTC--AGACAGGGCT
Ancestral sequences 9  CACCCCGCTTC--AGACAGGGCT
Olive baboon           CACCCCGCTTC--AGACAGGGCT
Ancestral sequences 10 CACCCCGCTTC--AGACAGGGCT
Gelada                 CACCCCGCTTC--AGACAGGGCT
Ancestral sequences 11 CACCCCGCTTC--AGACAGGGCT
Marmoset               CACCCCGCTTC--AGACAGGGCT
Ancestral sequences 12 CACTCCGCTTC--AGACAGGGCT
Mouse Lemur            CACTCTGCCTCCTCGGGTGGGAC
```

| RsID | Alleles | Effect | Trial Sample |
| --- | --- | --- | --- |
| rs2228485 | A G | Loneliness^3^  ASD^4^  Emotion recognition^5^ | 285  195 (Chinese) |
| rs237897 | G  A | ASD^6^  Altruism^7^  Lower self-reported betrayal levels^8^  Social connectedness^9^  Theory of Mind^10^ | 152  203  165  11.000  301 |
| rs11131149 | G  A | Theory of Mind, higher levels of social cognition^10^  Depressive mood^11^  Lower levels of social cognition^10^ | 350 children  493 (Japanese)  350 children |
| rs59190448 | A  G | Anxiety, stress and depression risk^12^ | 653 |
| rs13316193 | T  C | ASD^6^ Depressive mood^11^ Poor empathic communication^13^ Empathy^14^ Poor social skills^15^ Greater cooperation and comforting^16^ ADHD^17^  Face emotion recognition^18^ | 152  493 (Japanese)  120  101(Chinese)  112  422 (Chinese males)  276 ADHD patients  151 ADHD patients |
| rs9872310 | A  G | Altruism^7^  ASD^6^ | 203  152 |
| rs4686302 | C  T | Better perspective taking skills^14^  Face emotion recognition^19^  Social connectedness in men, opposite in women^9^ | 101(Chinese)  151 ADHD patients  >11000 individuals |
| rs237888 | C  T | IQ and VABS scores^6^  Altruism^7^  Greater impairment of ASD^20^  Methylation of CpG sites linked to abuse and psychiatric symptoms^21^ | 152  203  1002 ASD patients  393(African/American) |
| rs6770632 | A  G | Aggression^22^  VABS scores^6^ | 160 children  152 |
| rs237885 | G  T | Altruism^7^  ASD^23^  Schizophrenia^24^  Callous/unemotional traits^22^  Higher risk of aggression^25^ | 203  282 (Japanese)  145  160 children  488 cases, 488 control (Chinese) |
| rs1042778 |  | Lower levels of OT in plasma, diminished parental care^26^  Panic and aggressive behaviors^27^  Recovery from low maternal emotional warmth^28^  ASD^6^  Aggression^22^  Prosocial fund allocations in the Dictator Game^7^  Might lower transcription levels of  OTR^27^  Altruism, comforting behavior^14^  Positive emotions after training^29^ | 352  2341  152, 2333, 209  160  203  422 Chinese males  122 |
| rs237911 | A  G | ASD^4,23,30^ | 195 (Chinese),  282 (Japanese), 3941 |
| rs2254298 | G  A | Lower communication^13^  Variation in empathy^14^  Methylation at cg11589699 (increased depression and anxiety)^31^  Less sensitive parenting and lower plasma OXT^26^  Higher positive affect^24^  Lower scores in depressive temperament^11^  Higher levels of Retrospective Self-Report of Inhibition and Adult Separation Anxiety^32^  Smaller left amygdala^33^  ASD^4,23,30^  Lower levels of emotion recognition and resilience scores^34^  Increased amygdala volume^35^  Fewer social deficits in ADHD, more social deficits in ASD^18^  Lower serum OT in ASD patients^36^  Positive parenting behavior, physically controlling behavior^37^  Responsive to adversity^38^  High levels of physical aggression^39^ | 120  101 (Chinese)  393 (African American)  352  352  493 (Japanese)  93  211 men, 199 women  195 (Chinese), 282 (Japanese), 3941  264 (Korean)  55  341 (ASD), 276 (ADHD)  55 (ASD), 110 (controls)  157  302  197 (Chinese) |
| rs53576 | G  A | Bulimia Nerviosa^40^  Diminished stress^41^  Separation anxiety  Oxytocin sensitivity in social cooperation (increased in males, decreased in females)^42^  Weak social cognition in ADHD^15^  ASD^4,6^  Empathy^43,44^  Lower psychological resources^45^  Social connectedness (women)^9^ | 262 (Korean)  176  185  204  112  195 (Chinese), 152  50, 192  344  >11000 |
| rs2268490 | C  T | Altruism^7^  Vocal alterations under stress^46^  Stress-related vocal symptoms and higher cortisol levels^46^ | 203  657 Finnish twins  657 Finnish twins |
| rs2268493 | T | ASD^47–50^  Negative scores in social tasks in schizophrenia^51^  ADHD^52^  Depressive temperament^11^ | 417, 530(Caucasian), 527, 2.333  74  99  493 (Japanese) |
| rs237917 | T | Emotion recognition^53^ | 207 (Central European) |
| rs237889 | C  T | Utilitarian answers in dilemmas^54^  ASD^6^ | 228, 322  152 |

| **RsID** | **Alleles** | **Effect** | **Trial Sample** |
| --- | --- | --- | --- |
| rs1042615 | A | ASD^55^ | 205 (Finnish) |
| rs10784339 | G | Stress reactivity and substance addiction risk^56,57^ | 852, 2231 |
| rs11174811 | C | Substance addiction risk^56,57^  Higher anxiety levels^58^  Aggression^22^ | 852, 2231  1090 (German)  160 children |
| rs3021529 | G | Addiction^59^ | 1.050 |
| rs10877969 | A | ASD^60,61^ | 150 (Korean trios), 633 |

**Supplementary Table 7:** Literature review results for *OTR* identified Single Nucleotide Polymorphisms (SNPs). For each variant site, ‘RsID’ (Variant/SNP ID), the allele discussed in each study (‘Alleles’), a short note on the main findings (‘Effect’) and the ‘Trial sample’ used for each study are listed.

**Supplementary Table 8**: Literature review results for *VTR1A* identified Single Nucleotide Polymorphisms (SNPs). For each variant site, ‘RsID’ (Variant/SNP ID), the allele discussed in each study (‘Alleles’), a short note on the main findings (‘Effect’) and the ‘Trial sample’ used for each study are listed.

| **RsID** | **Alleles** | **Effect** | **Trial sample** |
| --- | --- | --- | --- |
| rs28676508 | T | Child onset aggression^62^ | 177 |
| rs28632197 | T | ASD^48^  Panic disorder^63^ | 207  186 (German) |
| rs33985287 | C | Protects against depressive moods^64^ | 464 (children) |

**Supplementary Table 9**: Literature review results for *VTR1B* identified Single Nucleotide Polymorphisms (SNPs). For each variant site, ‘RsID’ (Variant/SNP ID), the allele discussed in each study (‘Alleles’), a short note on the main findings (‘Effect’) and the ‘Trial sample’ used for each study are listed.
